# Critical periods when dopamine controls behavioral responding during Pavlovian learning

**DOI:** 10.1101/2022.02.28.482312

**Authors:** Merridee J. Lefner, Claire E. Stelly, Kaitlyn M. Fonzi, Hector Zurita, Matthew J. Wanat

## Abstract

**Rationale:** Learning the association between rewards and predictive cues is critical for appetitive behavioral responding. The mesolimbic dopamine system is thought to play an integral role in establishing these cue-reward associations. The dopamine response to cues can signal differences in reward value, though this emerges only after significant training. This suggests that the dopamine system may differentially regulate behavioral responding depending on the phase of training.

**Objectives:** The purpose of this study was to determine whether antagonizing dopamine receptors elicited different effects on behavior depending on the phase of training or the type of Pavlovian task.

**Methods:** Separate groups of male rats were trained on Pavlovian tasks in which distinct audio cues signaled either differences in reward size or differences in reward rate. The dopamine receptor antagonist flupenthixol was systemically administered prior to either the first ten sessions of training (acquisition phase) or the second ten sessions of training (expression phase) and we monitored the effect of these manipulations for an additional ten training sessions.

**Results:** We identified acute effects of dopamine receptor antagonism on conditioned responding, the latency to respond, and post-reward head entries in both Pavlovian tasks. Interestingly, dopamine receptor antagonism during the expression phase produced persistent deficits in behavioral responding only in rats trained on the reward size Pavlovian task.

**Conclusions:** Together, our results illustrate that dopamine’s control over behavior in Pavlovian tasks depends upon one’s prior training experience and the information signaled by the cues.

## Introduction

Learning to associate rewarding outcomes to the cues that predict them is a critical process for promoting efficient reward-seeking behavior. These cue-reward associations are regulated by the mesolimbic dopamine system (Phillips et al. 2007; Salamone and Correa 2012). Prior research illustrates that the delivery of a reward evokes a pronounced elevation in dopamine transmission during early Pavlovian training (Coddington and Dudman 2018; Day et al. 2007; Schultz et al. 1997). After the subject experiences multiple cue-reward pairings, the reward-evoked dopamine response decays and the cue-evoked dopamine response increases (Coddington and Dudman 2018; Day et al. 2007; Schultz et al. 1997). In well-trained animals, dopamine neurons respond to cues to encode differences in reward-related information including reward size (Gan et al. 2010; Lefner et al. 2022; Roesch et al. 2007; Tobler et al. 2005), reward probability (Fiorillo et al. 2003; Hart et al. 2015), and reward rate (Fonzi et al. 2017; Stelly et al. 2021). Our recent work suggests these value-related dopamine responses to cues emerge through a multi-step process. Specifically, cue-evoked dopamine release does not signal differences in reward size or reward rate during early Pavlovian training sessions, though these value-related dopamine signals emerge after extended training (Fonzi et al. 2017; Lefner et al. 2022; Stelly et al. 2021). However, it is unclear if dopamine’s control over behavioral responding in these Pavlovian tasks depends on (1) the experience with the task (early training vs well-trained) and (2) the information signaled by the reward-predictive cues (reward size or reward rate).

Dopamine signaling can influence the acquisition and expression of conditioned responding in appetitive Pavlovian tasks, though it should be noted that dopamine’s control over behavior can depend upon the design of the Pavlovian task (Di Ciano et al. 2001; Eyny and Horvitz 2003; Flagel et al. 2011; Fraser and Janak 2017; Horvitz 2001; Roughley and Killcross 2019; Saunders and Robinson 2012; Sculfort et al. 2016; Stelly et al. 2021; Stelly et al. 2020; Wassum et al. 2011). Prior studies illustrate that antagonizing dopamine receptors during early training sessions can acutely impair conditioned responding (Di Ciano et al. 2001; Flagel et al. 2011; Roughley and Killcross 2019; Sculfort et al. 2016; Stelly et al. 2021). Additionally, antagonizing dopamine receptors or optogenetic inhibition of dopamine neurons acutely suppresses conditioned responding in well-trained animals (Di Ciano et al. 2001; Fraser and Janak 2017; Heymann et al. 2020; Lee et al. 2020; Morrens et al. 2020; Saunders and Robinson 2012). These deficits in conditioned responding can persist beyond the sessions in which the dopamine receptor antagonist was administered, which illustrates dopamine signaling is critical for reward learning (Flagel et al. 2011; Roughley and Killcross 2019; Sculfort et al. 2016; Stelly et al. 2021; Stelly et al. 2020). However, it remains unclear if the behavioral alterations following dopamine receptor antagonism can be reversed with further training.

In this study, we trained separate groups of rats on two different Pavlovian tasks that elicit goal-tracking behavior. In the Pavlovian Reward Size task, two distinct audio cues signaled the delivery of either a small reward or a large reward (Lefner et al. 2022). In the Pavlovian Reward Rate task, two distinct audio cues both signaled the delivery of a small reward but differed in the time elapsed since the previous reward delivery (i.e. reward rate) (Fonzi et al. 2017; Stelly et al. 2021). The dopamine receptor antagonist flupenthixol was systemically administered during different phases of training to determine when the dopamine system regulates behavioral responding in these Pavlovian tasks. Rats received flupenthixol injections prior to either the first ten sessions of training (‘Acquisition phase’) or the second ten sessions of training (‘Expression phase’) and we monitored the effect of these manipulations for an additional ten training sessions. We quantified the effect of dopamine receptor antagonism on behavioral responding during the cue (e.g. conditioned responding and latency to respond) as well as the number of head entries following the reward delivery. Across Pavlovian tasks, flupenthixol treatment acutely impaired conditioned responding, the latency to respond, and post-reward head entries. However, dopamine receptor antagonism produced persistent deficits in behavioral responding exclusively during the Pavlovian Reward Size task. These findings highlight critical periods in which dopamine controls behavior in a manner that depends on the reward-related information signaled by the cues.

## Methods

### Subjects and surgery

All procedures were approved by the Institutional Animal Care and Use Committee at the University of Texas at San Antonio. Male Sprague-Dawley rats (Charles River, MA) weighing 300-350 g were pair-housed upon arrival and given ad libitum access to water and chow and maintained on a 12-hour light/dark cycle.

### Behavioral procedures

One week after arrival, rats were placed and maintained on mild food restriction (~15 g/day of standard lab chow) to target 90% free-feeding weight, allowing for an increase of 1.5% per week. Behavioral sessions were performed in chambers (Med Associates) that had grid floors, a house light, a food tray, a food tray light, and two auditory stimulus generators (4.5 kHz tone, and either 2.5 kHz tone or white noise). To familiarize rats with the chamber and food retrieval, rats underwent a single magazine training session in which 20 food pellets (45 mg, BioServ) were non-contingently delivered at a 90 ± 15 s variable interval. Rats then underwent 30 sessions (1/day) of Pavlovian training under the Pavlovian Reward Size or Pavlovian Reward Rate tasks as described previously (Fonzi et al. 2017; Lefner et al. 2022; Stelly et al. 2021).

Training sessions for the Pavlovian Reward Size task consisted of 50 trials where the termination of a 5 s audio CS (2.5 kHz tone or 4.5 kHz tone, counterbalanced across animals) resulted in the delivery of a single food pellet (US; Small Reward trial) and a separate CS resulted in the delivery of three food pellets (Large Reward trial). The food port light illuminated coinciding with reward delivery and remained on for 4.5 s. Each session contained 25 Small Reward trials and 25 Large Reward trials that were delivered in a pseudorandom order, with a 45 ± 5 s ITI between all trials. Training sessions for the Pavlovian Reward Rate task consisted of 50 trials where the termination of a 5 s audio cue (white noise or 4.5 kHz tone, counterbalanced across animals) resulted in the delivery of a single food pellet and illumination of the food port light for 4.5 s. Each session contained 25 High Rate trials in which a CS was presented after a 20 ± 5 s ITI, and 25 Low Rate trials in which a separate CS was presented after a 70 ± 5 s ITI. The High Rate and Low Rate trials were presented in a pseudorandom order. Conditioned responding was quantified as the change in the rate of head entries during the 5 s CS relative to the 5 s preceding the CS delivery (Fonzi et al. 2017; Lefner et al. 2022; Stelly et al. 2021). We also quantified the latency to initiate a head entry during the CS. For the post-US analysis, we calculated the number of head entries made during the 9 s following reward delivery. This 9 s post-reward window corresponds to 4.5 s in which the tray light in the food port was illuminated and an equivalent 4.5 s after the tray light turned off. Response vigor was calculated as the change in the rate of head entries between the first CS-evoked head entry and the end of the CS relative to the 5 s preceding the CS delivery (Stelly et al. 2020).

### Pharmacology

Flupenthixol dihydrochloride (Tocris) was dissolved in sterile 0.9% NaCl. Rats received i.p. injections of flupenthixol (225 ug/kg) or saline vehicle 1 hour prior to Pavlovian training sessions (Flagel et al. 2011). Flupenthixol injections were administered prior to sessions 1-10 (‘Acquisition phase’) or prior to sessions 11-20 (‘Expression phase’). No injections were administered prior to sessions 21-30. The group sizes are as follows: Pavlovian Reward Size groups: *n* = 8 control saline, *n* = 8 Acquisition phase flupenthixol, *n* = 8 Expression phase flupenthixol; Pavlovian Reward Size groups: *n* = 10 control saline, *n* = 10 Acquisition phase flupenthixol, *n* = 10 Expression phase flupenthixol.

### Data analysis

Statistical analyses were performed in GraphPad Prism 9. Behavioral responding was analyzed using a three-way mixed-effects model fit (restricted maximum likelihood method), using repeated measures where appropriate with treatment, session, and trial type as independent factors. The Geisser-Greenhouse correction was applied to address unequal variances between groups where applicable. If the mixed-effects model found a significant effect of treatment, a subsequent two-way ANOVA was performed for each trial type followed by a post-hoc Sidak’s test to identify differences between treatment groups on a session by session basis. Note that data are plotted separately by trial type for visual clarity in all figures. The full list of statistical analyses is presented in **Supplementary Information**.

## Results

Separate groups of rats were trained on two different Pavlovian tasks in which distinct audio cues signaled different reward values. The Reward Size Pavlovian task utilized one audio cue (CS) that signaled the delivery of a single sucrose pellet (US; Small Reward trial) and a distinct audio cue that signaled the delivery of three sucrose pellets (Large Reward trial, **Fig. 1A**). Each session contained 25 Small Reward trials and 25 Large Reward trials that were presented in a pseudorandom order. The Reward Rate Pavlovian task used two distinct 5 s audio CSs that both signaled the delivery of a single sucrose pellet, but the CSs differed in the time elapsed since the previous reward (Fonzi et al. 2017; Stelly et al. 2021). In High Rate trials the CS was presented 20 ± 5 s ITI following the previous reward delivery, and in Low Rate trials the CS was presented 70 ± 5 s following the previous reward delivery (**Fig. 2A**). Each session contained 25 Low Rate trials and 25 High Rate trials that were presented in a pseudorandom order. To determine how the dopamine system regulates behavioral responding across different phases of training, we systemically administered the non-selective dopamine receptor antagonist flupenthixol one hour prior to the first ten training sessions (‘Acquisition phase’) or the next ten training sessions (‘Expression phase’). Control groups were given saline injections for the first twenty sessions. No injections were administered during the final ten training sessions.

**Figure 1.**
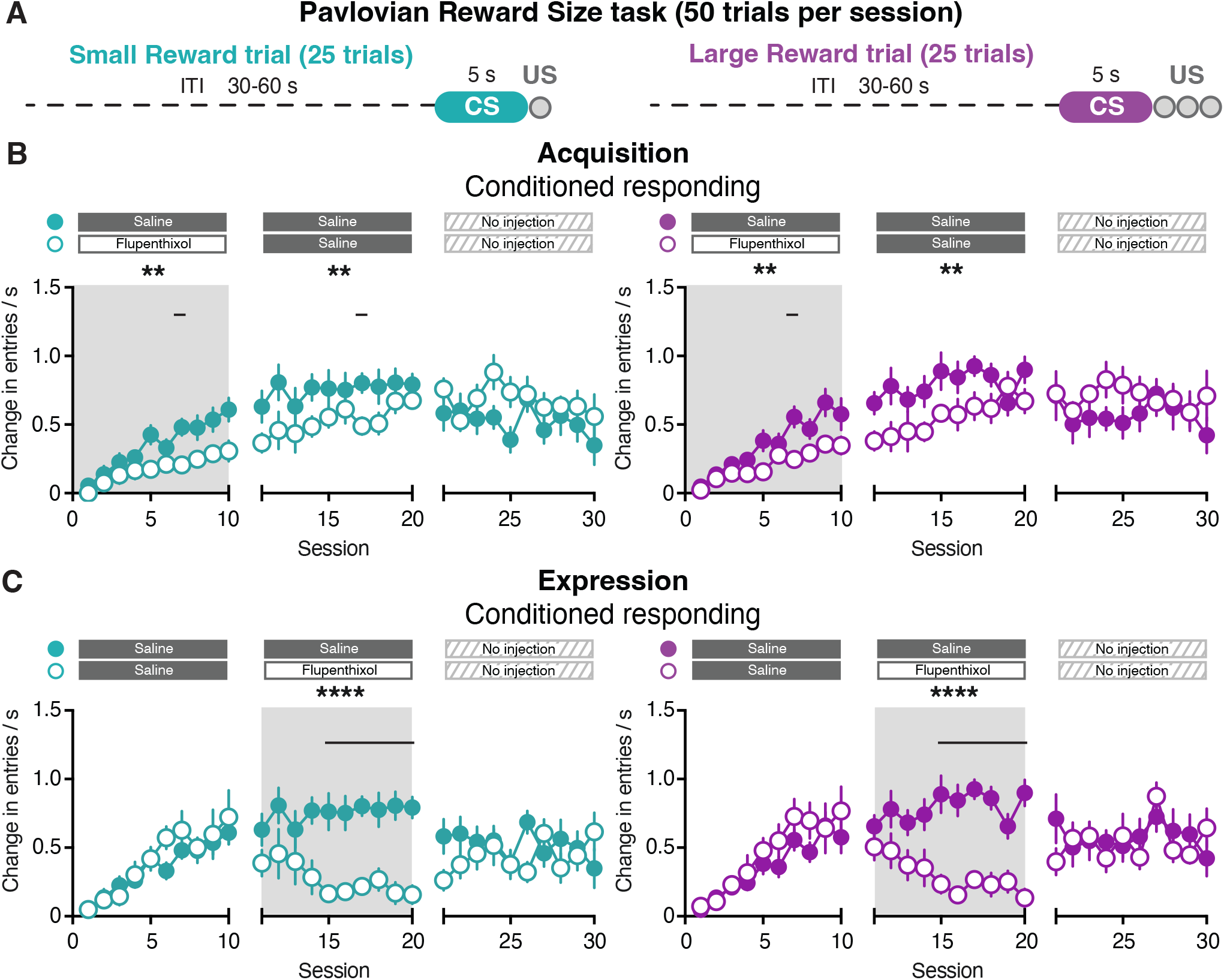
Conditioned responding during the Pavlovian Reward Size task. (A) Pavlovian Reward Size task design (B) Conditioned responding when flupenthixol is administered during the Acquisition phase (Sessions 1-10; open circles) compared to saline-injected control subjects (closed circles) for Small Reward trials (teal, left) and Large Reward trials (purple, right). Note that animals experienced both trials in the same session though the data are plotted separately by trial type for visual clarity. (C) Conditioned responding when flupenthixol is administered during the Expression phase (Sessions 11-20; open circles) compared to saline-injected control subjects (closed circles) for Small Reward trials (teal, left) and Large Reward trials (purple, right). Significant main effect of treatment: ** *p* < 0.01, **** *p* < 0.0001. Black lines above the graphs denote significant post-hoc effect of treatment on a given session (*p* < 0.05).

**Figure 2.**
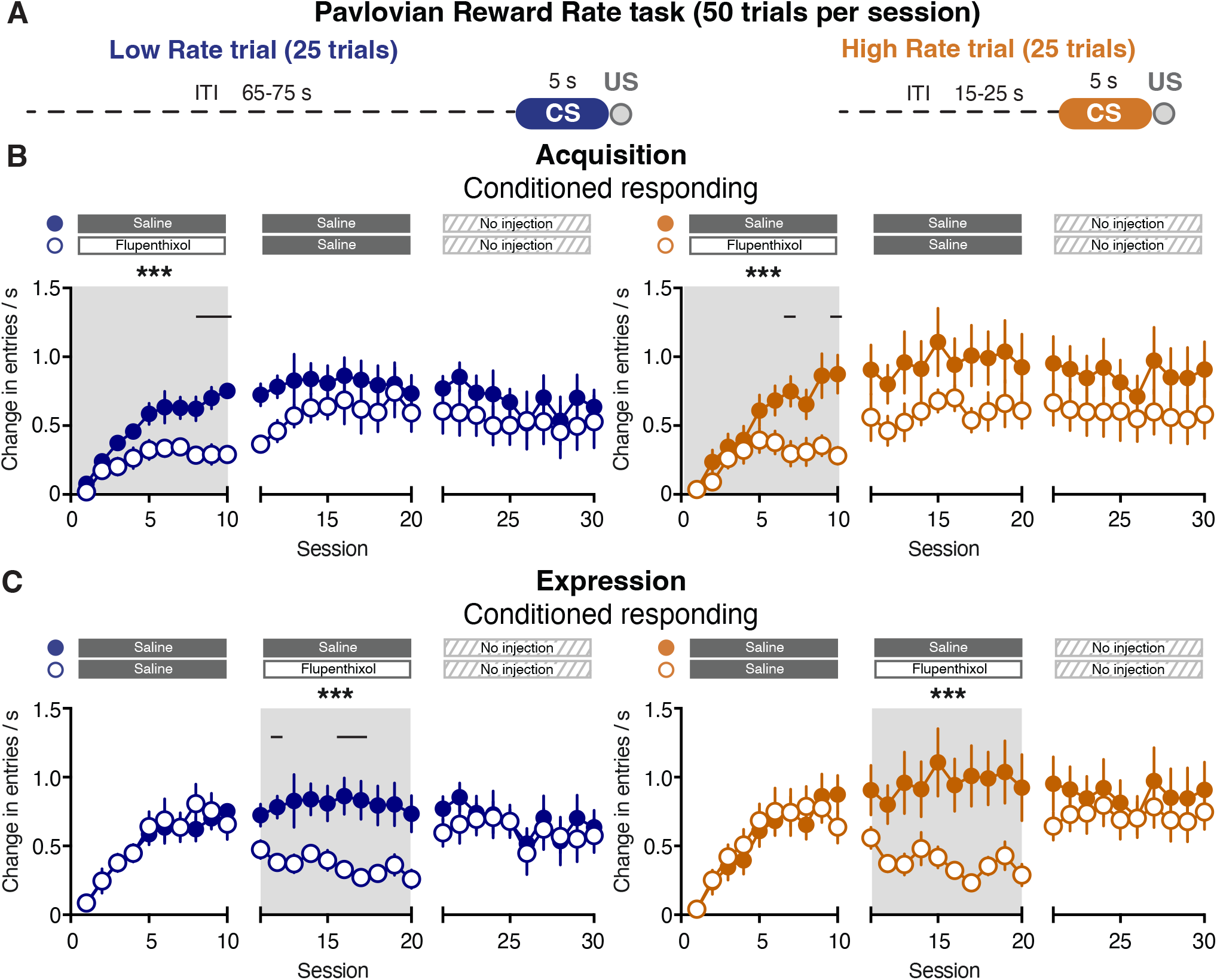
Conditioned responding during the Pavlovian Reward Rate task. (A) Pavlovian Reward Rate task design (B) Conditioned responding when flupenthixol is administered during the Acquisition phase (Sessions 1-10; open circles) compared to saline-injected control subjects (closed circles) for Low Rate trials (blue, left) and High Rate trials (orange, right). Note that animals experienced both trials in the same session though the data are plotted separately by trial type for visual clarity. (C) Conditioned responding when flupenthixol is administered during the Expression phase (Sessions 11-20; open circles) compared to saline-injected control subjects (closed circles) for Low Rate trials (blue, left) and High Rate trials (orange, right). Significant main effect of treatment: *** *p* < 0.001. Black lines above the graphs denote significant post-hoc effect of treatment on a given session (*p* < 0.05).

In the Pavlovian Reward Size task, there was no difference in conditioned responding between Small and Large Reward trials (Sessions 1-10 three-way mixed-effects analysis; reward size effect: *F*_(1, 14)_ = 2.52, *p* = 0.14; **Fig. 1B**), consistent with prior research (Lefner et al. 2022). Antagonizing dopamine receptors during the Acquisition phase impaired conditioned responding in both trial types (Sessions 1-10 three-way mixed-effects analysis; treatment effect: *F*_(1, 126)_ = 9.27, *p* = 0.003; session x treatment effect: *F*_(9, 126)_ = 2.53, *p* = 0.01; **Fig. 1B**). Conditioned responding remained lower in rats that had previously received flupenthixol during the Acquisition phase compared to control rats (Sessions 11-20; treatment effect: *F*_(1, 126)_ = 7.28, *p* = 0.008; **Fig. 1B**). However, there was no difference in conditioned responding between groups in the final ten sessions of training in which no injections were given (Sessions 21-30; treatment effect: *F*_(1, 126)_ = 1.26, *p* = 0.26; **Fig. 1B**).

In a separate group of rats that had already undergone ten training sessions we examined how antagonizing the dopamine system regulates conditioned responding during the Expression phase (**Fig. 1C**). Flupenthixol injections during the Expression phase acutely impaired conditioned responding during both Small and Large Reward trials (Sessions 11-20 three-way mixed-effects analysis; treatment effect: *F*_(1, 126)_ = 18.88, *p* < 0.0001; session x treatment effect: *F*_(9, 126)_ = 2.74, *p* = 0.006; **Fig. 1C**). There was no effect of treatment during the following ten sessions in which no injections were given (Sessions 21-30; treatment effect: *F*_(1, 126)_ = 0.25, *p* = 0.62; session x treatment effect: *F*_(9, 126)_ = 3.06, *p* = 0.002; **Fig. 1C**). Together this indicates that antagonizing dopamine receptors during early training sessions produced both acute and prolonged decreases in conditioned responding in rats trained on the Pavlovian Reward Size task. However, flupenthixol treatment in well-trained animals on this task produced only an acute decrease in conditioned responding.

In rats trained on the Pavlovian Reward Rate task, there was no difference in conditioned responding between Low and High Rate trials (Sessions 1-10 three-way mixed-effects analysis; reward rate effect: *F*_(1, 18)_ = 0.29, *p* = 0.60; **Fig. 2B**), consistent with prior studies (Fonzi et al. 2017; Stelly et al. 2021). Similar to the Pavlovian Reward Size task, antagonizing dopamine receptors during the Acquisition phase impaired conditioned responding in both trial types (Sessions 1-10 three-way mixed-effects analysis; treatment effect: *F*_(1, 162)_ = 12.78, *p* = 0.0005; session x treatment effect: *F*_(9, 162)_ = 6.29, *p* < 0.0001; **Fig. 2B**). In contrast to the Pavlovian Reward Size task, rats did not display a prolonged decrease in conditioned responding during the next ten sessions of training (Sessions 11-20; treatment effect: *F*_(1, 162)_ = 3.06, *p* = 0.08; **Fig. 2B**). However when examining the first session following flupenthixol treatment, conditioned responding was reduced relative to controls (Session 11 two-way mixed-effects analysis; treatment effect: *F*_(1, 14)_ = 6.11, *p* = 0.02; **Fig. 2B**). Therefore in the Pavlovian Reward Rate task, the deficits in conditioned responding during the Acquisition phase can rapidly reverse when flupenthixol is no longer administered. Antagonizing dopamine receptors during the Expression phase diminished conditioned responding in both Low and High Rate trials (Sessions 11-20 three-way mixed-effects analysis; treatment effect: *F*_(1, 162)_ = 11.90, *p* = 0.0007; **Fig. 2C**), though there was no difference between treatment groups in the next ten training sessions following flupenthixol administration (Sessions 21-30; treatment effect: *F*_(1, 162)_ = 0.47, *p* = 0.49; **Fig. 2C**). These results illustrate that antagonizing dopamine receptors in rats trained on the Pavlovian Reward Rate task produced acute and rapidly reversible deficits on conditioned responding.

We additionally examined the effects of dopamine receptor antagonism on the latency to enter the food port following the onset of the CS (**Fig. 3**). In rats trained on the Pavlovian Reward Size task, antagonizing dopamine receptors during the Acquisition phase increased the latency to respond in both Small and Large Reward trials (Sessions 1-10 three-way mixed-effects analysis; treatment effect: *F*_(1, 126)_ = 8.88, *p* = 0.004; **Fig. 3A**). However there were no prolonged effects on the latency to respond in the next ten sessions (Sessions 11-20; treatment effect: *F*_(1, 122)_ = 1.07, *p* = 0.30; **Fig. 3A**). Antagonizing dopamine receptors during the Expression phase increased the latency to respond in both trial types (Sessions 11-20 three-way mixed-effects analysis; treatment effect: *F*_(1, 126)_ = 55.73, *p* < 0.0001; session x treatment effect: *F*_(9, 126)_ = 4.55, *p* < 0.0001; **Fig. 3B**). Additionally, the latency to respond remained elevated in the last set of sessions where no injections were administered (Sessions 21-30; treatment effect: *F*_(1, 126)_ = 7.13, *p* = 0.009; session x treatment effect: *F*_(9, 126)_ = 2.26, *p* = 0.02; **Fig. 3B**). Together, these results illustrate that perturbing dopamine signaling in well-trained animals produces sustained deficits in the latency to respond, without affecting conditioned responding in rats trained on the Pavlovian Reward Size task (**Fig. 1C, Fig. 3B**).

**Figure 3.**
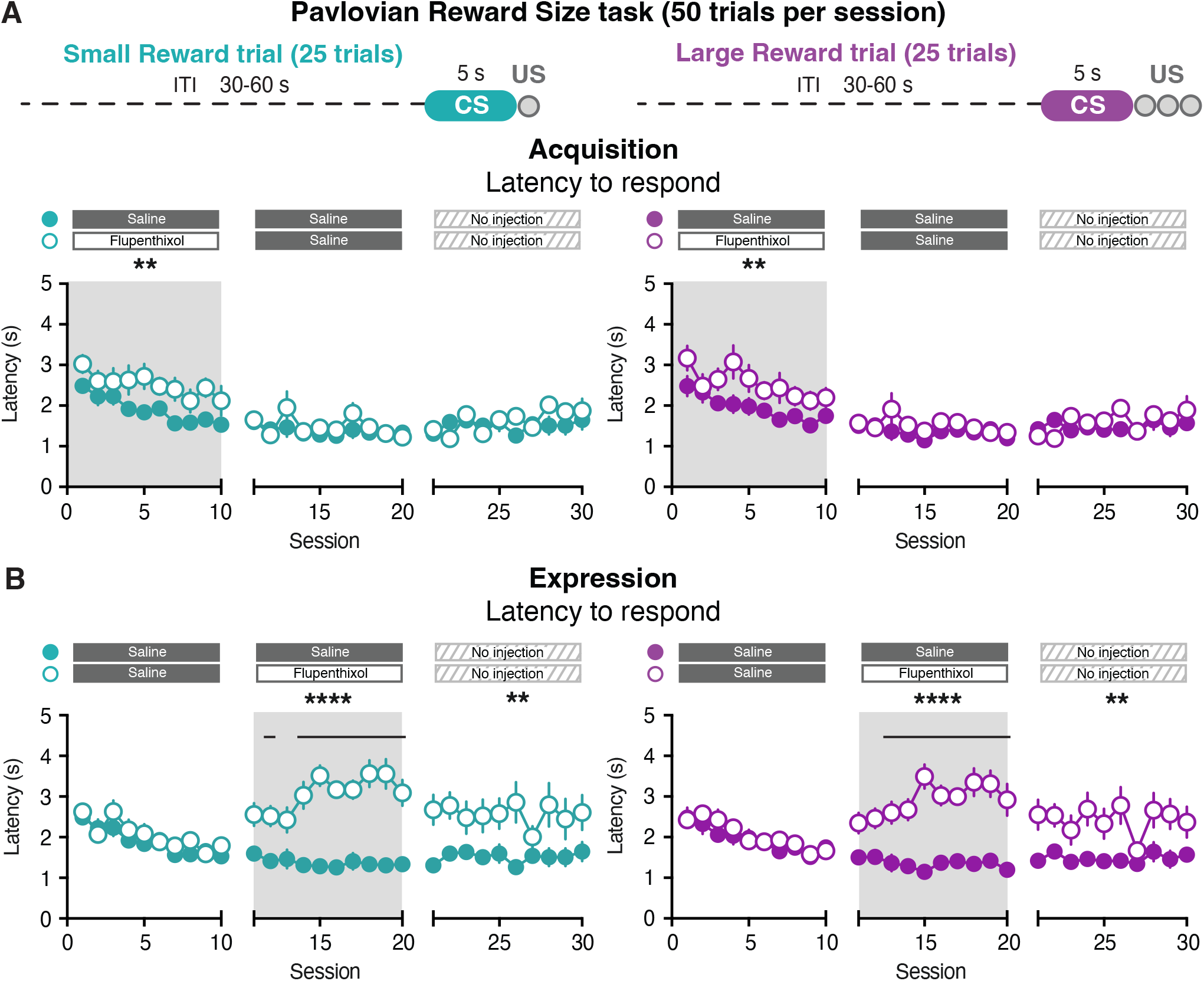
Latency to respond during the Pavlovian Reward Size task.(A) Latency to respond when flupenthixol is administered during the Acquisition phase for Small and Large Reward trials. (B) Latency to respond when flupenthixol is administered during the Expression phase for Small and Large Reward trials. Significant main effect of treatment: ** *p* < 0.01, **** *p* < 0.0001. Black lines above the graphs denote significant post-hoc effect of treatment on a given session (*p* < 0.05).

In rats trained on the Pavlovian Reward Rate task, antagonizing dopamine receptors during the Acquisition phase also increased the latency to respond in both Low and High Rate trials (Sessions 1-10 three-way mixed-effects analysis; treatment effect: *F*_(1, 162)_ = 5.81, *p* = 0.02; **Fig. 4A**), though there was no difference between groups in the following ten sessions (Sessions 11-20; treatment effect: *F*_(1, 162)_ = 1.22, *p* = 0.27; **Fig. 4A**). Similarly, antagonizing dopamine receptors during the Expression phase increased the latency to respond in both trial types (Sessions 11-20; treatment effect: *F*_(1, 162)_ = 15.09, *p* = 0.0001; session x treatment effect: *F*_(9, 162)_ = 2.42, *p* < 0.01; **Fig. 4B**), with no prolonged effects in the following sessions (Sessions 21-30; treatment effect: *F*_(1, 162)_ = 0.79, *p* = 0.37; **Fig. 4B**). Collectively, these results demonstrate that the influence of the dopamine system on cue-evoked behavioral responding depends on the phase of training as well as whether the cues denote differences in reward size (**Figs. 1,3**) or reward rate (**Figs. 2,4**).

**Figure 4.**
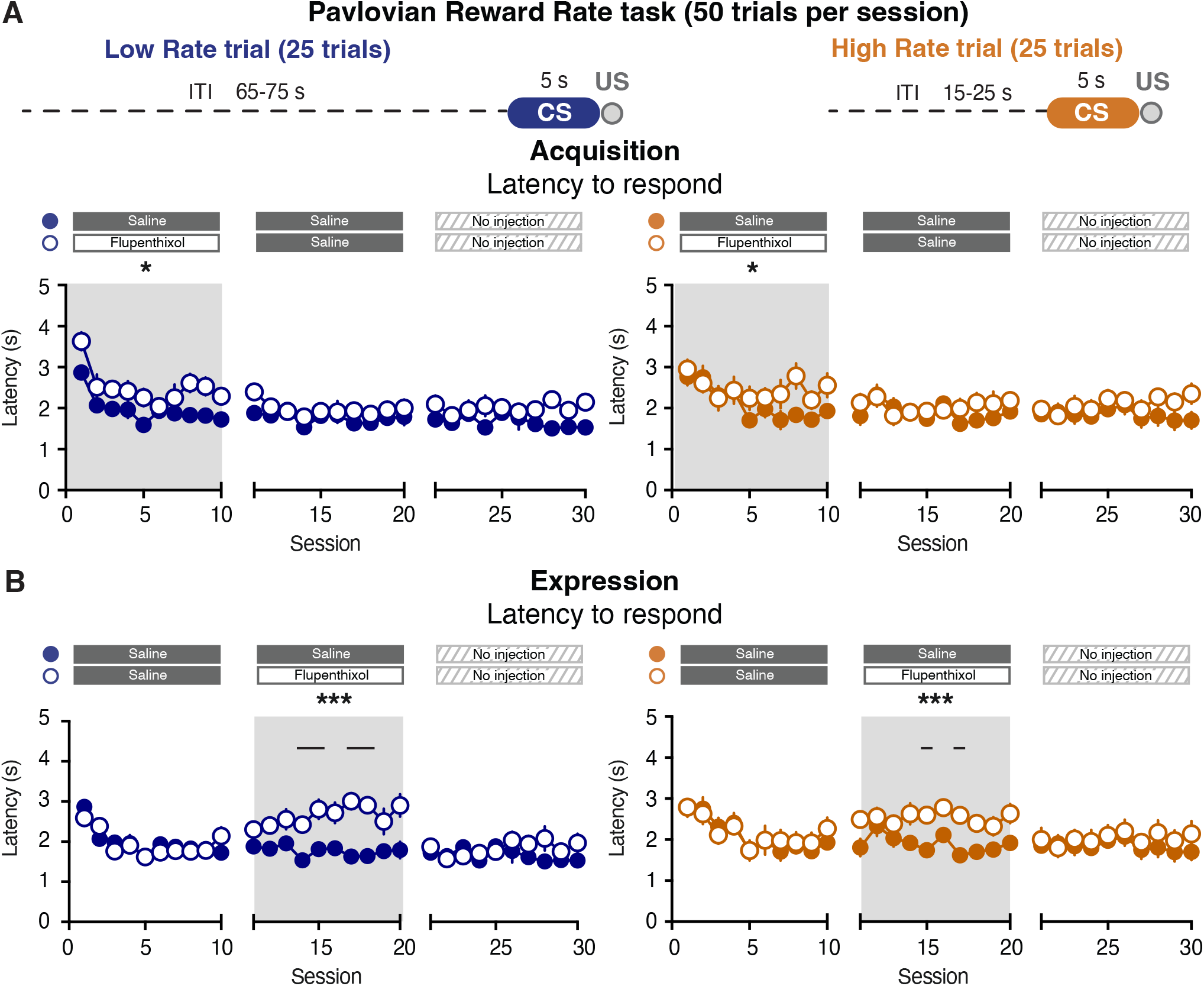
Latency to respond during the Pavlovian Reward Rate task (A) Latency to respond when flupenthixol is administered during the Acquisition phase for Low and High Rate trials. (B) Latency to respond when flupenthixol is administered during the Expression phase for Low and High Rate trials. Significant main effect of treatment: * *p* < 0.05, *** *p* < 0.001. Black lines above the graphs denote significant post-hoc effect of treatment on a given session (*p* < 0.05).

We next examined behavioral responses following reward delivery by quantifying the number of head entries occurring in the 9 s after the termination of the CS (Post US; **Fig. 5A**). Rats perform a greater number of head entries following the delivery of a large reward relative to the delivery of a small reward (Lefner et al. 2022). Flupenthixol administration during the Acquisition phase decreased Post US head entries for both trial types in the Pavlovian Reward Size task (Sessions 1-10 three-way mixed-effects analysis; treatment effect: *F*_(1, 126)_ = 16.73, *p* < 0.0001; session x treatment effect: *F*_(9, 126)_ = 2.11, *p* = 0.03; three-way interaction effect: *F*_(9, 126)_ = 2.82, *p* = 0.005; **Fig. 5B**). This effect was also observed in the first session following flupenthixol treatment (Session 11 two-way mixed-effects analysis; treatment effect: *F*_(1, 14)_ = 5.48, *p* = 0.03; Sessions 11-20 three-way mixed-effects analysis; treatment effect: *F*_(1, 126)_ = 3.35, *p* = 0.07; **Fig. 5B**). Furthermore, the impairments in Post US head entries were also evident during the final ten sessions in which no injections were administered (Sessions 21-30; treatment effect: *F*_(1, 126)_ = 4.47, *p* = 0.04; **Fig. 5B**). Antagonizing dopamine receptors during the Expression phase decreased Post US head entries in both trial types (Sessions 11-20 three-way mixed-effects analysis; treatment effect: *F*_(1, 126)_ = 5.23, *p* = 0.02; **Fig. 5C**). This effect was also observed in the ten sessions following flupenthixol treatment (Sessions 21-30; treatment effect: *F*_(1, 126)_ = 4.61, *p* = 0.03; **Fig. 5C**). These results suggest that dopamine receptor antagonism produces both acute and persistent deficits on Post US head entries in rats trained on the Pavlovian Reward Size task.

**Figure 5.**
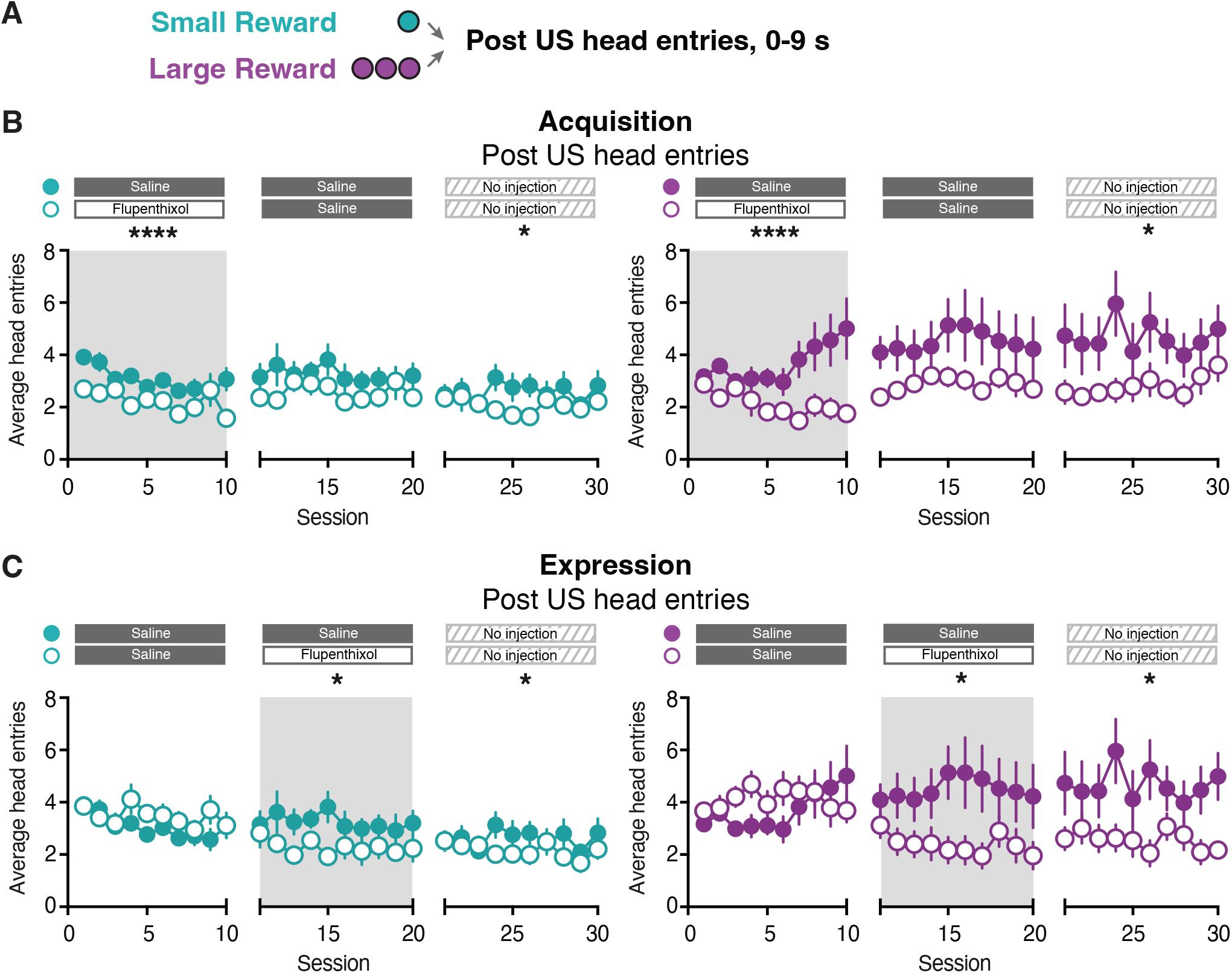
Head entries following reward delivery during the Pavlovian Reward Size task. (A) Post US time window (B) Post US head entries when flupenthixol is administered during the Acquisition phase for Small and Large Reward trials. (B) Post US head entries when flupenthixol is administered during the Expression phase for Small and Large Reward trials. Significant main effect of treatment: * *p* < 0.05, **** *p* < 0.0001.

In the Pavlovian Reward Rate task, flupenthixol administration during the Acquisition phase diminished Post US head entries during both Low and High Rate trials (Sessions 1-10 three-way mixed-effects analysis; treatment effect: *F*_(1, 162)_ = 4.29, *p* = 0.04; **Fig. 6A-B**), with no effect on the following ten sessions (Sessions 11-20; treatment effect: *F*_(1, 162)_ = 2.28, *p* = 0.13; **Fig. 6B**). In contrast, dopamine receptor antagonism during the Expression phase did not produce acute or prolonged effects on Post US head entries (Sessions 11-20; three-way mixed-effects analysis; treatment effect: *F*_(1, 162)_ = 0.64, *p* = 0.42; Sessions 21-30; treatment effect: *F*_(1, 162)_ = 0.38, *p* = 0.54; **Fig. 6C**). These results collectively highlight that the influence of the dopamine system on Post US responding depends on the phase of Pavlovian training as well as the size of the delivered reward (**Figs. 5-6**).

**Figure 6.**
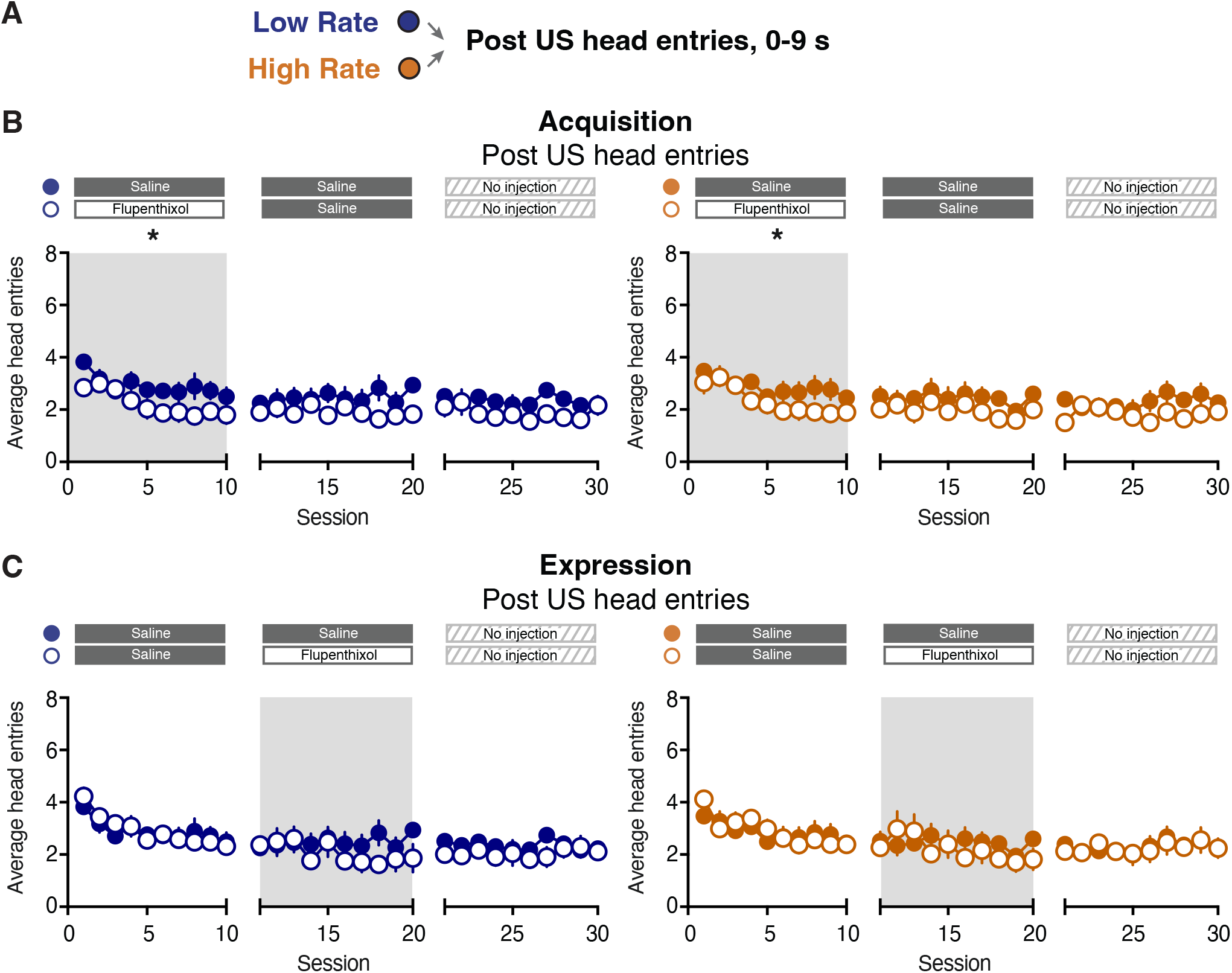
Head entries following reward delivery during the Pavlovian Reward Rate task. (A) Post US time window (B) Post US head entries when flupenthixol is administered during the Acquisition phase for Low and High Rate trials. (B) Post US head entries when flupenthixol is administered during the Expression phase for Low and High Rate trials. Significant main effect of treatment: * *p* < 0.05.

Prolonged deficits in behavioral responses following dopamine receptor antagonism during early training sessions provides evidence for dopamine’s role in Pavlovian learning (Flagel et al. 2011; Roughley and Killcross 2019; Sculfort et al. 2016; Stelly et al. 2021). However, our current results illustrate that flupenthixol treatment during the Expression phase in the Pavlovian Reward Size task produces sustained deficits in the latency to respond and in Post US head entries (**Fig. 3C, Fig. 5C**). These flupenthixol-elicited behavioral alterations could arise from a sustained suppression in the number of head entries performed by the subject. We therefore examined how dopamine receptor antagonism impacted the total number of head entries across sessions. Flupenthixol treatment during the Acquisition phase acutely decreased the total number of head entries in rats trained on the Pavlovian Reward Size task (Sessions 1-10 two-way mixed-effects model; treatment effect: *F*_(1, 14)_ = 16.24, *p* = 0.001; **Fig. 7A**), with no sustained effect during the following sessions (Sessions 11-20; treatment effect: *F*_(1, 14)_ = 2.54, *p* = 0.13; **Fig. 7A**). In contrast, flupenthixol treatment during the Expression phase acutely decreased the total number of head entries (Sessions 11-20 two-way mixed-effects model; treatment effect: *F*_(1, 14)_ = 27.09, *p* = 0.001; **Fig. 7A**) and produced sustained deficits in the following sessions without drug injections (Sessions 21-30; treatment effect: *F*_(1, 14)_ = 6.89, *p* = 0.02; **Fig. 7A**). While dopamine receptor antagonism reduced the total number of head entries in the last ten training sessions of the Pavlovian Reward Size task, this did not impact the CS-evoked change in head entries (i.e. conditioned responding; **Fig. 1C**), or the CS-evoked change in response vigor (**Supplementary Fig. 1**). In the Pavlovian Reward Rate task, sustained effects on the total number of head entries were not observed when rats received flupenthixol injections during the Acquisition phase (Sessions 1-10 two-way mixed-effects model; treatment effect: *F*_(1, 18)_ = 9.47, *p* = 0.007; Sessions 11-20; treatment effect: *F*_(1, 18)_ = 3.02, p = 0.10; **Fig. 7B**) or the Expression phase (Sessions 11-20 two-way mixed-effects model; treatment effect: *F*_(1, 18)_ = 11.85, *p* = 0.003; Sessions 21-30; treatment effect: *F*_(1, 18)_ = 0.03, *p* = 0.86; **Fig. 7B**). A summary of the observed effects of dopamine receptor antagonism on cue- and reward-evoked behavioral responses for both Pavlovian tasks can be found in **Fig. 8**.

**Figure 7.**
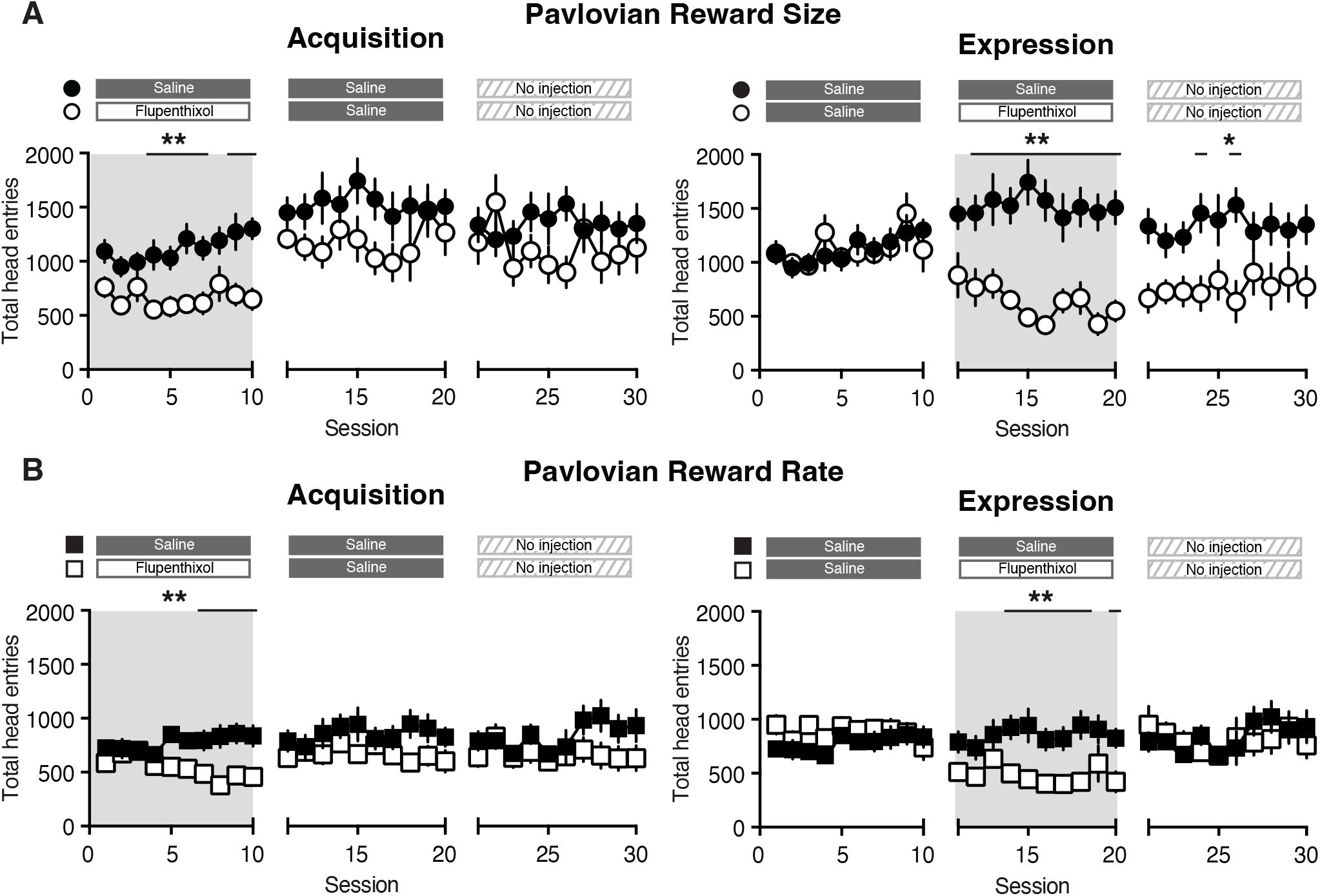
Total head entries (A) Left; Total head entries in the Pavlovian Reward Size task when flupenthixol is administered during the Acquisition phase (open circles) compared to saline-injected control subjects (closed circles). Right; Total head entries when flupenthixol is administered during the Expression phase. (B) Left; Total head entries in the Pavlovian Reward Rate task when flupenthixol is administered during the Acquisition phase (open squares) compared to saline-injected control subjects (closed squares). Right; Total head entries when flupenthixol is administered during the Expression phase. Significant main effect of treatment: * *p* < 0.05, ** *p* < 0.01. Black lines above the graphs denote significant post-hoc effect of treatment on a given session (*p* < 0.05).

**Figure 8.**
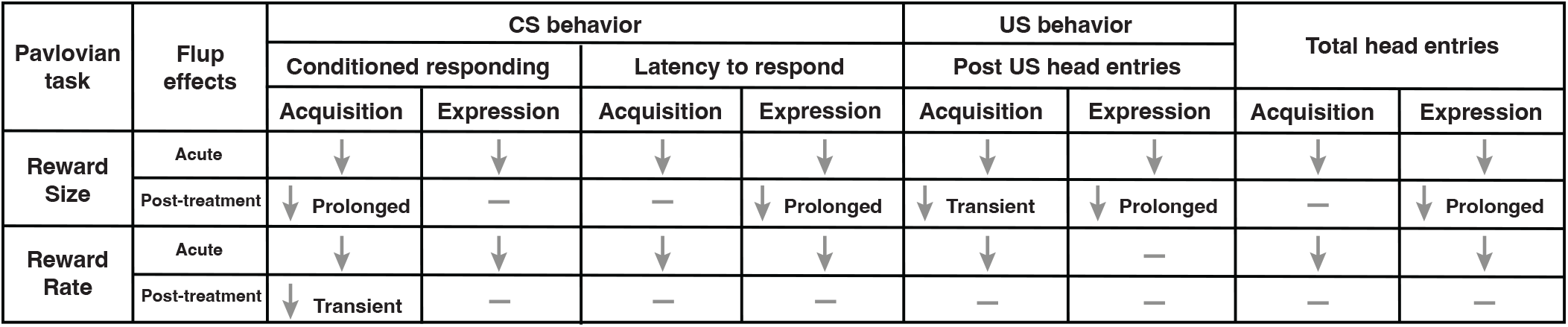
Summary table demonstrating the main effects of treatment administered during the Acquisition phase or the Expression phase on cue- and reward-evoked behaviors during both Pavlovian tasks. Acute effects of flupenthixol on behavior are represented by arrows; sustained post-treatment effects of flupenthixol on behavior are indicated with text.

## Discussion

Our findings demonstrate that perturbations of the dopamine system can alter behavioral responding throughout Pavlovian training. Prior research illustrates that dopamine receptor antagonism or optogenetic inhibition of dopamine neurons acutely suppresses conditioned responding (Di Ciano et al. 2001; Fraser and Janak 2017; Heymann et al. 2020; Lee et al. 2020; Morrens et al. 2020; Saunders and Robinson 2012). In support, we also found acute effects of dopamine receptor antagonism on both cue- and reward-evoked behavior across Pavlovian tasks. However, we note that the post-reward head entries during the Expression phase of the Pavlovian Reward Rate task were not impacted by flupenthixol injections. This illustrates that the array of behavioral responses exhibited during Pavlovian conditioning are not uniformly regulated by dopamine signaling.

Many lines of evidence highlight that dopamine is involved with learning in Pavlovian conditioning tasks (Flagel et al. 2011; Roughley and Killcross 2019; Sculfort et al. 2016; Stelly et al. 2021). In support, we find that flupenthixol treatment during early training sessions produced a transient decrease in conditioned responding in the sessions following drug treatment in the Pavlovian Reward Rate task. However, this same manipulation resulted in a prolonged decrease in conditioned responding in the Pavlovian Reward Size task. Paralleling these effects on conditioned responding, flupenthixol treatment produced a sustained decrease in post-reward head entries and total head entries in the Pavlovian Reward Size task, but no prolonged effects on post-reward head entries or total head entries in the Pavlovian Reward Rate task. We note that it can be a challenge to disentangle the effect of dopamine receptor antagonism on learning versus performance, especially when distinct behaviors can reach asymptotic performance on different days. In this study, we operationally referred to injections during the first ten sessions as the ‘Acquisition phase’ and injections during the second ten sessions as the ‘Expression phase’. However, we acknowledge these training windows may not define when learning has ended for a given behavior. As such, we focused on highlighting the acute and post-injection effects of flupenthixol treatment. Regardless, future studies will be needed to parse out which sessions of training are most critical for dopamine receptor antagonism to elicit prolonged behavioral deficits for a given behavior.

It is unclear what neurobiological mechanism accounts for the differential effect of flupenthixol treatment on behavioral responding between the Reward Size and Reward Rate Pavlovian tasks. However, we note that subjects earn an average of two food pellets per trial in the Pavlovian Reward Size task whereas subjects earn an average of one food pellet per trial in the Pavlovian Reward Rate task. We speculate that suppressing dopamine signaling may functionally produce a negative prediction error when the reward is delivered. Therefore, antagonizing dopamine receptors when the expected average reward size is high may then produce longer lasting behavioral effects relative to when the average reward size is low.

The Pavlovian tasks in our study used audio cues which elicit goal-tracking behavior. Prior studies have found that dopamine receptor antagonism can acutely disrupt goal-tracking behavior (Flagel et al. 2011; Roughley and Killcross 2019; Sculfort et al. 2016; Stelly et al. 2021; Wassum et al. 2011). Our results extend on these findings as we demonstrate both acute and sustained effects of flupenthixol treatment on conditioned responding in the Pavlovian Reward Size task. However, it is important to acknowledge that the sustained effects of dopamine receptor antagonism depends on the information signaled by the cues as well as the modality of cue utilized. For example in a task where subjects can perform both sign- and goal-tracking responses, prolonged effects of dopamine receptor antagonism are not observed on goal-tracking behavior (Flagel et al. 2011). Therefore, one must exercise caution when comparing the results between Pavlovian studies without considering what types of behavioral responding is elicited by the cues.

Our experimental approach utilized systemic injections of flupenthixol, which is a non-selective dopamine receptor antagonist. Prior research illustrates that administration of either a D1 or D2 receptor antagonist decreases goal-tracking behavior (Fraser et al. 2016; Khoo et al. 2021; Lopez et al. 2015; Roughley and Killcross 2019), but see (Eyny and Horvitz 2003; Horvitz 2001). As such, future studies are necessary to determine the role of D1- and D2-type receptors in the acquisition and expression of behavioral responding in the Reward Size and Reward Rate Pavlovian tasks. By performing systemic drug injections, it remains unclear where in the brain dopamine is acting to regulate behavioral responses in these Pavlovian tasks. Prior studies have implicated a role for dopamine signaling in the ventral striatum, as local injections of flupenthixol into the nucleus accumbens core decreased conditioned responding during sessions in which the antagonist was administered (Di Ciano et al. 2001; Fraser and Janak 2017; Saunders and Robinson 2012; Stelly et al. 2021). Prolonged deficits on conditioned responding are observed for a least one session following local injections of flupenthixol into the nucleus accumbens core, the nucleus accumbens shell, or the ventral lateral striatum (Stelly et al. 2021; Stelly et al. 2020). Therefore further experiments are needed to determine where dopamine is mediating the acute and prolonged effects on behavioral responses during different Pavlovian tasks.

We found no differences in conditioned responding between trial types in either Pavlovian task. However, recordings of dopamine release in animals trained on these tasks demonstrate that reward-evoked dopamine release signals differences in reward size during early training sessions and that cue-evoked dopamine release signals differences in reward rate and reward size in well-trained subjects (Fonzi et al. 2017; Lefner et al. 2022). These findings collectively illustrate that a difference in cue-evoked dopamine release between trials is not driving a corresponding difference in conditioned responding. Rather our prior results demonstrate that when the dopamine response to a given cue changes in well-trained subjects, the animal then exhibits a corresponding update in conditioned responding toward that cue (Fonzi et al. 2017). Future studies are needed to determine if updates in cue-evoked dopamine release similarly controls changes in the latency to respond or post-reward head entries as well as if these behavioral effects are observed across sexes. In sum, our data demonstrates that dopamine’s control over responding during Pavlovian conditioning depends on (i) what behavior is being studied, (ii) prior training experience, and (iii) the information signaled by the cues.

