## Supplementary Information for "Critical periods when dopamine controls behavioral responding during Pavlovian learning"

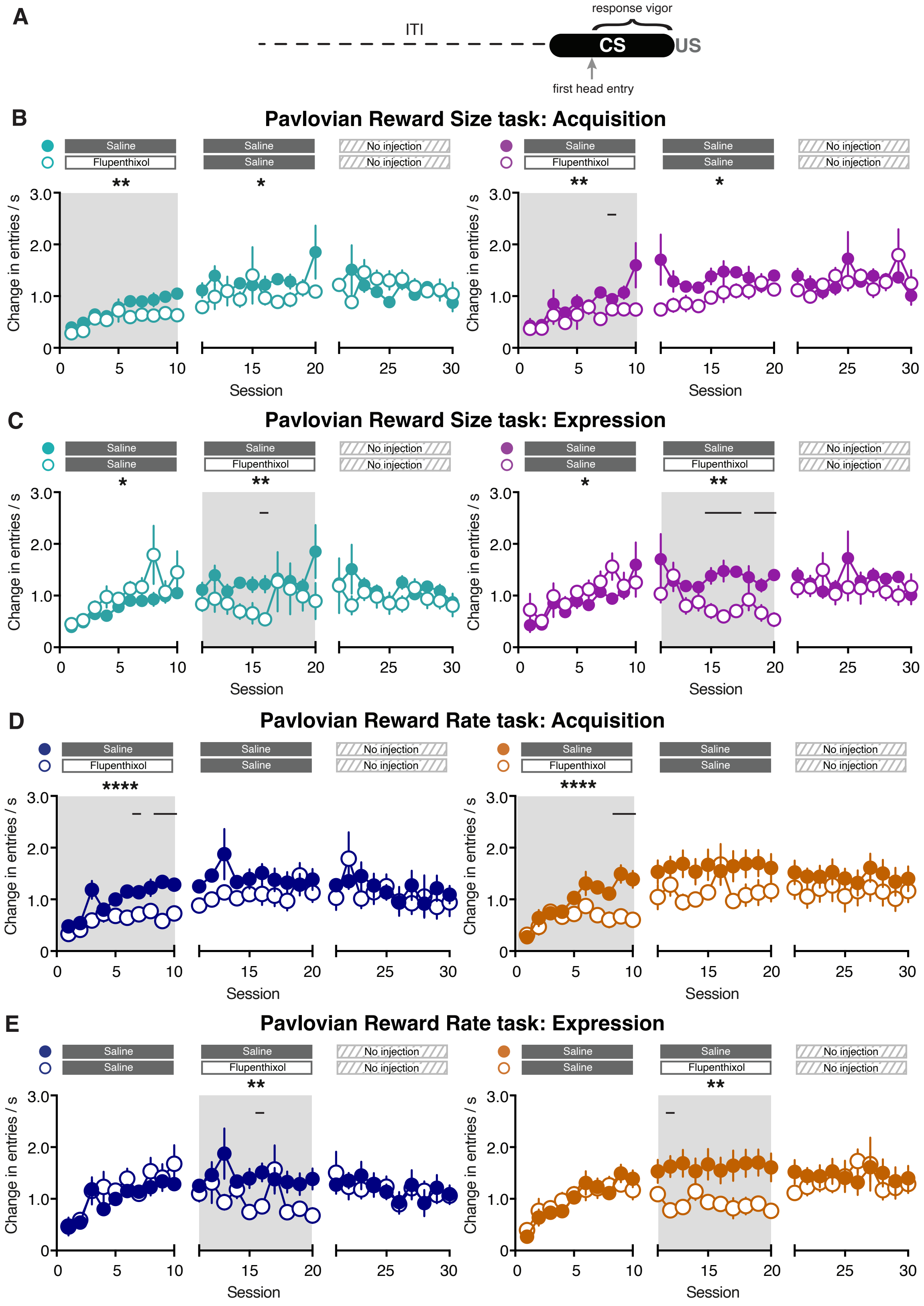


Supplementary Figure 1

**Supplementary Figure 1** Response vigor (A) Calculation of response vigor. (B) Response vigor when flupenthixol is administered during the Acquisition phase for Small and Large Reward trials in the Pavlovian Reward Size task (Sessions 1-10 three-way mixed-effects analysis; session effect: $F_{(3.19, 44.67)} = 9.83$, $p < 0.0001$; treatment effect: $F_{(1, 126)} = 8.79$, $p = 0.004$; reward size effect: $F_{(1, 14)} = 3.78$, $p = 0.07$; session x treatment effect: $F_{(9, 126)} = 1.74$, $p = 0.09$; session x reward size effect: $F_{(2.88, 40.28)} = 0.58$, $p = 0.62$; treatment x reward size effect: $F_{(1, 126)} = 0.62$, $p = 0.43$; three-way interaction effect: $F_{(9, 126)} = 0.67$, $p = 0.73$; Sessions 11-20 three-way mixed-effects analysis; session effect: $F_{(2.76, 38.65)} = 0.97$, $p = 0.97$; treatment effect: $F_{(1, 126)} = 4.99$, $p = 0.03$; reward size effect: $F_{(1, 14)} = 0.06$, $p = 0.81$; session x treatment effect: $F_{(9, 126)} = 1.00$, $p = 0.44$; session x reward size effect: $F_{(2.96, 41.42)} = 1.14$, $p = 0.34$; treatment x reward size effect: $F_{(1, 126)} = 0.68$, $p = 0.41$; three-way interaction effect: $F_{(9, 126)} = 1.23$, $p = 0.28$; Sessions 21-30 three-way mixed-effects analysis; session effect: $F_{(3.77, 52.77)} = 0.84$, $p = 0.50$; treatment effect: $F_{(1, 126)} = 0.03$, $p = 0.86$; reward size effect: $F_{(1, 14)} = 6.45$, $p = 0.02$; session x treatment effect: $F_{(9, 126)} = 1.27$, $p = 0.26$; session x reward size effect: $F_{(3.72, 52.13)} = 1.75$, $p = 0.16$; treatment x reward size effect: $F_{(1, 126)} = 0.62$, $p = 0.43$; three-way interaction effect: $F_{(9, 126)} = 1.53$, $p = 0.14$). (C) Response vigor when flupenthixol is administered during the Expression phase for Small and Large Reward trials in the Pavlovian Reward Size task (Sessions 1-10 three-way mixed-effects analysis; session effect: $F_{(3.13, 43.88)} = 9.93$, $p < 0.0001$; treatment effect: $F_{(1, 126)} = 3.99$, $p < 0.05$; reward size effect: $F_{(1, 14)} = 0.43$, $p = 0.52$; session x treatment effect: $F_{(9, 126)} = 1.17$, $p = 0.32$; session x reward size effect: $F_{(3.10, 43.34)} = 0.46$, $p = 0.71$; treatment x reward size effect: $F_{(1, 126)} = 0.17$, $p = 0.68$; three-way interaction effect: $F_{(9, 126)} = 0.68$, $p = 0.73$; Sessions 11-20 three-way mixed-effects analysis; session effect: $F_{(4.12, 57.65)} = 0.90$, $p = 0.47$; treatment effect: $F_{(1, 126)} = 11.33$, $p = 0.001$; reward size effect: $F_{(1, 14)} = 0.01$, $p = 0.91$; session x treatment effect: $F_{(9, 126)} = 0.95$, $p = 0.49$; session x reward size effect: $F_{(3.66, 51.26)} = 1.41$, $p = 0.25$; treatment x reward size effect: $F_{(1, 126)} = 1.12$, $p = 0.29$; three-way interaction effect: $F_{(9, 126)} = 0.87$, $p = 0.55$; Sessions 21-30 three-way mixed-effects analysis; session effect: $F_{(4.12, 55.67)} = 0.52$, $p = 0.73$; treatment effect: $F_{(1, 126)} = 1.17$, $p = 0.28$; reward size effect: $F_{(1, 14)} = 6.49$, $p = 0.02$; session x treatment effect: $F_{(9, 126)} = 0.56$, $p = 0.83$; session x reward size effect: $F_{(4.46, 62.45)} = 1.01$, $p = 0.41$; treatment x reward size effect: $F_{(1, 126)} = 0.08$, $p = 0.77$; three-way interaction effect: $F_{(9, 126)} = 1.50$, $p = 0.15$). (D) Response vigor when flupenthixol is administered during the Acquisition phase for Low and High Rate trials in the Pavlovian Reward Rate task (Sessions 1-10 three-way mixed-effects analysis; session effect: $F_{(3.94, 70.90)} = 15.40$, $p < 0.0001$; treatment effect: $F_{(1, 162)} = 17.04$, $p < 0.0001$; reward rate effect: $F_{(1, 18)} = 0.0001$, $p = 0.99$; session x treatment effect: $F_{(9, 162)} = 4.59$, $p < 0.0001$; session x reward rate effect: $F_{(5.29, 95.13)} = 1.45$, $p = 0.21$; treatment x reward size effect: $F_{(1, 162)} = 0.10$, $p = 0.75$; three-way interaction effect: $F_{(9, 162)} = 1.55$, $p = 0.14$; Sessions 11-20 three-way mixed-effects analysis; session effect: $F_{(3.07, 55.28)} = 1.03$, $p = 0.39$; treatment effect: $F_{(1, 162)} = 3.79$, $p = 0.05$; reward rate effect: $F_{(1, 18)} = 1.53$, $p = 0.23$; session x treatment effect: $F_{(9, 162)} = 0.90$, $p = 0.53$; session x reward rate effect: $F_{(4.50, 81.02)} = 1.11$, $p = 0.36$; treatment x reward rate effect: $F_{(1, 162)} = 0.51$, $p = 0.48$; three-way interaction effect: $F_{(9, 162)} = 1.60$, $p = 0.12$; Sessions 21-30 three-way mixed-effects analysis; session effect: $F_{(3.83, 68.94)} = 1.40$, $p = 0.25$; treatment effect: $F_{(1, 162)} = 0.48$, $p = 0.49$; reward rate effect: $F_{(1, 18)} = 2.99$, $p = 0.10$; session x treatment effect: $F_{(9, 162)} = 0.62$, $p = 0.78$; session x reward rate effect: $F_{(4.09, 73.69)} = 1.23$, $p = 0.27$; treatment x reward size effect: $F_{(1, 162)} = 1.23$, $p = 0.27$; three-way interaction effect: $F_{(9, 162)} = 1.37$, $p = 0.20$). (E) Response vigor when flupenthixol is administered during the Expression phase for Low and High Reward trials in the Pavlovian Reward Rate task. (Sessions 1-10 three-way mixed-effects analysis; session effect: $F_{(3.36, 60.51)} = 19.08$, $p < 0.0001$; treatment effect: $F_{(1, 162)} = 0.75$, $p = 0.39$; reward rate effect: $F_{(1, 18)} = 0.79$, $p = 0.38$; session x treatment effect: $F_{(9, 162)} = 0.60$, $p = 0.80$; session x reward rate effect: $F_{(4.36, 78.52)} = 1.18$, $p = 0.33$; treatment x reward size effect: $F_{(1, 162)} = 0.50$, $p = 0.48$; three-way interaction effect: $F_{(9, 162)} = 0.68$, $p = 0.73$; Sessions 11-20 three-way mixed-effects analysis; session effect: $F_{(4.04, 72.72)} = 0.87$, $p = 0.49$; treatment effect: $F_{(1, 162)} = 9.53$, $p = 0.002$; reward rate effect: $F_{(1, 18)} = 0.34$, $p = 0.56$; session x treatment effect: $F_{(9, 162)} = 1.52$, $p = 0.15$; session x reward rate effect: $F_{(3.60, 64.74)} = 1.42$, $p = 0.24$; treatment x reward rate effect: $F_{(1, 162)} = 2.16$, $p = 0.14$; three-way interaction effect: $F_{(9, 162)} = 1.14$, $p = 0.34$; Sessions 21-30 three-way mixed-effects analysis; session effect: $F_{(2.87, 51.65)} = 1.13$, $p = 0.35$; treatment effect: $F_{(1, 162)} = 0.06$, $p = 0.80$; reward rate effect: $F_{(1, 18)} = 3.36$, $p = 0.08$; session x treatment effect: $F_{(9, 162)} = 0.55$, $p = 0.84$; session x reward rate effect: $F_{(4.75, 85.57)} = 2.23$, $p = 0.06$; treatment x reward size effect: $F_{(1, 162)} = 0.17$, $p = 0.68$; three-way interaction effect: $F_{(9, 162)} = 1.54$, $p = 0.14$). Significant main effect of treatment: * $p < 0.05$, ** $p < 0.01$, **** $p < 0.0001$. Black lines above the graphs denote significant post-hoc effect of treatment on a given session ($p < 0.05$).

### Supplementary Table

| Table 1: Fig 1 – Conditioned responding – Pavlovian Reward Size task | | | |
|---|---|---|---|
| **Panel B – Acquisition: Pavlovian Reward Size task** | | | |
| Sessions 1-10<br>Three-way mixed-effects model | Session<br>$F_{(4.11, 57.53)} = 20.59$, $\boldsymbol{p < 0.0001}$ | Treatment<br>$F_{(1, 126)} = 9.27$, $\boldsymbol{p = 0.003}$ | Reward size<br>$F_{(1, 14)} = 2.52$, $p = 0.14$ |
| Session x Treatment<br>$F_{(9, 126)} = 2.53$, $\boldsymbol{p = 0.01}$ | Session x Reward size<br>$F_{(4.44, 62.14)} = 1.76$, $p = 0.14$ | Treatment x Reward size<br>$F_{(1, 126)} = 0.55$, $p = 0.46$ | Three-way interaction<br>$F_{(9, 126)} = 0.47$, $p = 0.89$ |
| Sessions 11-20<br>Three-way mixed-effects model | Session<br>$F_{(3.42, 47.82)} = 2.41$, $p = 0.07$ | Treatment<br>$F_{(1, 126)} = 7.28$, $\boldsymbol{p = 0.008}$ | Reward size<br>$F_{(1, 14)} = 5.08$, $\boldsymbol{p = 0.04}$ |
| Session x Treatment<br>$F_{(9, 126)} = 0.81$, $p = 0.60$ | Session x Reward size<br>$F_{(3.02, 42.33)} = 1.22$, $p = 0.32$ | Treatment x Reward size<br>$F_{(1, 126)} = 0.08$, $p = 0.78$ | Three-way interaction<br>$F_{(9, 126)} = 1.19$, $p = 0.31$ |
| Sessions 21-30<br>Three-way mixed-effects model | Session<br>$F_{(4.52, 63.20)} = 1.62$, $p = 0.17$ | Treatment<br>$F_{(1, 126)} = 1.26$, $p = 0.26$ | Reward size<br>$F_{(1, 14)} = 2.66$, $p = 0.13$ |
| Session x Treatment<br>$F_{(9, 126)} = 1.31$, $p = 0.24$ | Session x Reward size<br>$F_{(2.79, 39.03)} = 1.27$, $p = 0.30$ | Treatment x Reward size<br>$F_{(1, 126)} = 0.29$, $p = 0.59$ | Three-way interaction<br>$F_{(9, 126)} = 1.32$, $p = 0.23$ |
| Session 11<br>Two-way mixed-effects model | Reward size<br>$F_{(1, 14)} = 0.25$, $p = 0.63$ | Treatment<br>$F_{(1, 14)} = 5.63$, $\boldsymbol{p = 0.03}$ | Two-way interaction<br>$F_{(1, 14)} = 0.02$, $p = 0.90$ |
| **Panel C – Expression: Pavlovian Reward Size task** | | | |
| Sessions 1-10<br>Three-way mixed-effects model | Session<br>$F_{(2.92, 40.89)} = 20.54$, $\boldsymbol{p < 0.0001}$ | Treatment<br>$F_{(1, 126)} = 0.65$, $p = 0.42$ | Reward size<br>$F_{(1, 14)} = 3.56$, $p = 0.08$ |
| Session x Treatment<br>$F_{(9, 126)} = 0.78$, $p = 0.63$ | Session x Reward size<br>$F_{(3.51, 49.08)} = 1.27$, $p = 0.29$ | Treatment x Reward size<br>$F_{(1, 126)} = 1.63$, $p = 0.20$ | Three-way interaction<br>$F_{(9, 126)} = 1.34$, $p = 0.22$ |
| Sessions 11-20<br>Three-way mixed-effects model | Session<br>$F_{(3.46, 48.38)} = 0.73$, $p = 0.56$ | Treatment<br>$F_{(1, 126)} = 18.88$, $\boldsymbol{p < 0.0001}$ | Reward size<br>$F_{(1, 14)} = 0.10$, $p = 0.75$ |
| Session x Treatment<br>$F_{(9, 126)} = 2.74$, $\boldsymbol{p = 0.006}$ | Session x Reward size<br>$F_{(2.91, 40.75)} = 0.81$, $p = 0.49$ | Treatment x Reward size<br>$F_{(1, 126)} = 0.002$, $p = 0.96$ | Three-way interaction<br>$F_{(9, 126)} = 1.80$, $p = 0.08$ |
| Sessions 21-30<br>Three-way mixed-effects model | Session<br>$F_{(4.45, 62.22)} = 1.23$, $p = 0.31$ | Treatment<br>$F_{(1, 126)} = 0.25$, $p = 0.62$ | Reward size<br>$F_{(1, 14)} = 9.76$, $\boldsymbol{p = 0.008}$ |
| Session x Treatment<br>$F_{(9, 126)} = 3.06$, $\boldsymbol{p = 0.002}$ | Session x Reward size<br>$F_{(4.31, 60.27)} = 2.53$, $\boldsymbol{p < 0.05}$ | Treatment x Reward size<br>$F_{(1, 126)} = 1.18$, $p = 0.28$ | Three-way interaction<br>$F_{(9, 126)} = 1.26$, $p = 0.26$ |
| Session 21<br>Two-way mixed-effects model | Reward size<br>$F_{(1, 14)} = 6.95$, $\boldsymbol{p = 0.02}$ | Treatment<br>$F_{(1, 14)} = 3.50$, $p = 0.08$ | Two-way interaction<br>$F_{(1, 14)} = 0.01$, $p = 0.95$ |

| Table 2: Fig 2 – Conditioned responding – Pavlovian Reward Rate task | | | |
|---|---|---|---|
| **Panel B – Acquisition: Pavlovian Reward Rate task** | | | |
| Sessions 1-10<br>Three-way mixed-effects model | Session<br>$F_{(2.14, 38.52)} = 27.54$, $\boldsymbol{p < 0.0001}$ | Treatment<br>$F_{(1, 162)} = 12.78$, $\boldsymbol{p = 0.0005}$ | Reward rate<br>$F_{(1, 18)} = 0.29$, $p = 0.60$ |
| Session x Treatment<br>$F_{(9, 162)} = 6.29$, $\boldsymbol{p < 0.0001}$ | Session x Reward rate<br>$F_{(4.52, 81.39)} = 1.22$, $p = 0.31$ | Treatment x Reward rate<br>$F_{(1, 162)} = 0.03$, $p = 0.86$ | Three-way interaction<br>$F_{(9, 162)} = 1.58$, $p = 0.12$ |
| Sessions 11-20<br>Three-way mixed-effects model | Session<br>$F_{(2.16, 38.94)} = 1.46$, $p = 0.24$ | Treatment<br>$F_{(1, 162)} = 3.06$, $p = 0.08$ | Reward rate<br>$F_{(1, 18)} = 0.95$, $p = 0.34$ |
| Session x Treatment<br>$F_{(9, 162)} = 0.27$, $p = 0.98$ | Session x Reward rate<br>$F_{(3.60, 64.72)} = 1.19$, $p = 0.32$ | Treatment x Reward rate<br>$F_{(1, 162)} = 0.86$, $p = 0.36$ | Three-way interaction<br>$F_{(9, 162)} = 1.01$, $p = 0.44$ |
| Sessions 21-30<br>Three-way mixed-effects model | Session<br>$F_{(2.11, 38.01)} = 1.83$, $p = 0.17$ | Treatment<br>$F_{(1, 162)} = 1.09$, $p = 0.30$ | Reward rate<br>$F_{(1, 18)} = 2.34$, $p = 0.14$ |
| Session x Treatment<br>$F_{(9, 162)} = 0.58$, $p = 0.81$ | Session x Reward rate<br>$F_{(4.79, 86.21)} = 1.28$, $p = 0.28$ | Treatment x Reward rate<br>$F_{(1, 162)} = 0.64$, $p = 0.43$ | Three-way interaction<br>$F_{(9, 162)} = 0.61$, $p = 0.79$ |
| Session 11<br>Two-way mixed-effects model | Reward rate<br>$F_{(1, 18)} = 2.33$, $p = 0.14$ | Treatment<br>$F_{(1, 14)} = 6.11$, $\boldsymbol{p = 0.02}$ | Two-way interaction<br>$F_{(1, 14)} = 0.003$, $p = 0.96$ |
| **Panel C – Expression: Pavlovian Reward Rate task** | | | |
| Sessions 1-10<br>Three-way mixed-effects model | Session<br>$F_{(1.47, 26.43)} = 42.26$, $\boldsymbol{p < 0.0001}$ | Treatment<br>$F_{(1, 162)} = 0.06$, $p = 0.80$ | Reward rate<br>$F_{(1, 18)} = 0.15$, $p = 0.71$ |

| | | | |
|---|---|---|---|
| Session x Treatment<br>$F_{(9, 162)} = 1.22$, $p = 0.29$ | Session x Reward rate<br>$F_{(3.91, 70.36)} = 0.70$, $p = 0.59$ | Treatment x Reward rate<br>$F_{(1, 162)} = 0.004$, $p = 0.95$ | Three-way interaction<br>$F_{(9, 162)} = 0.55$, $p = 0.84$ |
| Sessions 11-20<br>Three-way mixed-effects model | Session<br>$F_{(2.27, 40.89)} = 1.06$, $p = 0.36$ | Treatment<br>$F_{(1, 162)} = 11.90$, $\mathbf{p = 0.0007}$ | Reward rate<br>$F_{(1, 18)} = 1.64$, $p = 0.22$ |
| Session x Treatment<br>$F_{(9, 162)} = 1.58$, $p = 0.13$ | Session x Reward rate<br>$F_{(5.14, 92.47)} = 1.13$, $p = 0.35$ | Treatment x Reward rate<br>$F_{(1, 162)} = 0.90$, $p = 0.34$ | Three-way interaction<br>$F_{(9, 162)} = 0.60$, $p = 0.80$ |
| Sessions 21-30<br>Three-way mixed-effects model | Session<br>$F_{(1.96, 35.26)} = 1.89$, $p = 0.17$ | Treatment<br>$F_{(1, 162)} = 0.47$, $p = 0.49$ | Reward rate<br>$F_{(1, 18)} = 2.44$, $p = 0.14$ |
| Session x Treatment<br>$F_{(9, 162)} = 0.55$, $p = 0.83$ | Session x Reward rate<br>$F_{(4.56, 82.07)} = 1.95$, $p = 0.10$ | Treatment x Reward rate<br>$F_{(1, 162)} = 0.17$, $p = 0.68$ | Three-way interaction<br>$F_{(9, 162)} = 0.68$, $p = 0.73$ |
| Session 21<br>Two-way mixed-effects model | Reward rate<br>$F_{(1, 14)} = 1.34$, $p = 0.26$ | Treatment<br>$F_{(1, 14)} = 2.54$, $p = 0.13$ | Two-way interaction<br>$F_{(1, 14)} = 0.45$, $p = 0.51$ |

| **Table 3: Fig 3 – Latency to respond – Pavlovian Reward Size task** | | | |
|---|---|---|---|
| **Panel A – Acquisition: Pavlovian Reward Size task** | | | |
| Sessions 1-10<br>Three-way mixed-effects model | Session<br>$F_{(3.90, 54.53)} = 5.00$, $\mathbf{p = 0.002}$ | Treatment<br>$F_{(1, 126)} = 8.88$, $\mathbf{p = 0.004}$ | Reward size<br>$F_{(1, 14)} = 1.02$, $p = 0.33$ |
| Session x Treatment<br>$F_{(9, 126)} = 0.51$, $p = 0.86$ | Session x Reward size<br>$F_{(4.69, 65.62)} = 1.58$, $p = 0.18$ | Treatment x Reward size<br>$F_{(1, 126)} = 0.06$, $p = 0.81$ | Three-way interaction<br>$F_{(9, 126)} = 0.72$, $p = 0.69$ |
| Sessions 11-20<br>Three-way mixed-effects model | Session<br>$F_{(2.47, 34.63)} = 2.34$, $p = 0.10$ | Treatment<br>$F_{(1, 122)} = 1.07$, $p = 0.30$ | Reward size<br>$F_{(1, 14)} = 0.01$, $p = 0.92$ |
| Session x Treatment<br>$F_{(9, 122)} = 1.03$, $p = 0.42$ | Session x Reward size<br>$F_{(3.01, 40.74)} = 1.66$, $p = 0.19$ | Treatment x Reward size<br>$F_{(1, 122)} = 0.48$, $p = 0.49$ | Three-way interaction<br>$F_{(9, 122)} = 0.84$, $p = 0.60$ |
| Sessions 21-30<br>Three-way mixed-effects model | Session<br>$F_{(3.44, 48.17)} = 2.85$, $\mathbf{p = 0.04}$ | Treatment<br>$F_{(1, 126)} = 0.51$, $p = 0.48$ | Reward size<br>$F_{(1, 14)} = 0.77$, $p = 0.40$ |
| Session x Treatment<br>$F_{(9, 126)} = 2.57$, $\mathbf{p < 0.01}$ | Session x Reward size<br>$F_{(3.12, 43.70)} = 1.38$, $p = 0.26$ | Treatment x Reward size<br>$F_{(1, 126)} = 0.01$, $p = 0.95$ | Three-way interaction<br>$F_{(9, 126)} = 1.33$, $p = 0.23$ |
| Session 11<br>Two-way mixed-effects model | Reward size<br>$F_{(1, 14)} = 1.72$, $p = 0.21$ | Treatment<br>$F_{(1, 14)} = 0.19$, $p = 0.67$ | Two-way interaction<br>$F_{(1, 14)} = 0.01$, $p = 0.94$ |
| **Panel B – Expression: Pavlovian Reward Size task** | | | |
| Sessions 1-10<br>Three-way mixed-effects model | Session<br>$F_{(4.00, 55.96)} = 13.57$, $\mathbf{p < 0.0001}$ | Treatment<br>$F_{(1, 126)} = 0.68$, $p = 0.41$ | Reward size<br>$F_{(1, 14)} = 0.23$, $p = 0.64$ |
| Session x Treatment<br>$F_{(9, 126)} = 0.60$, $p = 0.80$ | Session x Reward size<br>$F_{(4.87, 68.10)} = 1.58$, $p = 0.18$ | Treatment x Reward size<br>$F_{(1, 126)} = 0.66$, $p = 0.42$ | Three-way interaction<br>$F_{(9, 126)} = 1.16$, $p = 0.33$ |
| Sessions 11-20<br>Three-way mixed-effects model | Session<br>$F_{(3.32, 46.50)} = 2.32$, $p = 0.08$ | Treatment<br>$F_{(1, 126)} = 55.73$, $\mathbf{p < 0.0001}$ | Reward size<br>$F_{(1, 14)} = 5.01$, $\mathbf{p = 0.04}$ |
| Session x Treatment<br>$F_{(9, 126)} = 4.55$, $\mathbf{p < 0.0001}$ | Session x Reward size<br>$F_{(4.14, 57.96)} = 0.41$, $p = 0.66$ | Treatment x Reward size<br>$F_{(1, 126)} = 3.18$, $p = 0.08$ | Three-way interaction<br>$F_{(9, 126)} = 1.01$, $p = 0.43$ |
| Sessions 21-30<br>Three-way mixed-effects model | Session<br>$F_{(3.38, 47.34)} = 2.33$, $p = 0.08$ | Treatment<br>$F_{(1, 126)} = 7.13$, $\mathbf{p = 0.009}$ | Reward size<br>$F_{(1, 14)} = 3.81$, $p = 0.07$ |
| Session x Treatment<br>$F_{(9, 126)} = 2.26$, $\mathbf{p = 0.02}$ | Session x Reward size<br>$F_{(4.62, 64.64)} = 1.86$, $p = 0.12$ | Treatment x Reward size<br>$F_{(1, 126)} = 1.36$, $p = 0.25$ | Three-way interaction<br>$F_{(9, 126)} = 0.83$, $p = 0.59$ |
| Session 21<br>Two-way mixed-effects model | Reward size<br>$F_{(1, 14)} = 0.004$, $p = 0.95$ | Treatment<br>$F_{(1, 14)} = 10.28$, $\mathbf{p = 0.006}$ | Two-way interaction<br>$F_{(1, 14)} = 1.92$, $p = 0.19$ |

| **Table 4: Fig 4 – Latency to respond – Pavlovian Reward Rate task** | | | |
|---|---|---|---|
| **Panel A – Acquisition: Pavlovian Reward Rate task** | | | |
| Sessions 1-10<br>Three-way mixed-effects model | Session<br>$F_{(3.18, 57.19)} = 9.80$, $\mathbf{p < 0.0001}$ | Treatment<br>$F_{(1, 162)} = 5.81$, $\mathbf{p = 0.02}$ | Reward rate<br>$F_{(1, 18)} = 0.17$, $p = 0.68$ |
| Session x Treatment<br>$F_{(9, 162)} = 1.33$, $p = 0.22$ | Session x Reward rate<br>$F_{(4.42, 79.54)} = 3.89$, $\mathbf{p = 0.005}$ | Treatment x Reward rate<br>$F_{(1, 162)} = 0.52$, $p = 0.47$ | Three-way interaction<br>$F_{(9, 162)} = 2.11$, $\mathbf{p = 0.03}$ |
| Sessions 11-20<br>Three-way mixed-effects model | Session<br>$F_{(3.13, 56.39)} = 1.67$, $p = 0.18$ | Treatment<br>$F_{(1, 162)} = 1.22$, $p = 0.27$ | Reward rate<br>$F_{(1, 18)} = 1.10$, $p = 0.31$ |

| | | | |
|---|---|---|---|
| Session x Treatment $F_{(9, 162)} = 1.07$, $p = 0.39$ | Session x Reward rate $F_{(5.15, 92.76)} = 2.07$, $p = 0.07$ | Treatment x Reward rate $F_{(1, 162)} = 0.08$, $p = 0.77$ | Three-way interaction $F_{(9, 162)} = 0.75$, $p = 0.66$ |
| Sessions 21-30 Three-way mixed-effects model | Session $F_{(2.92, 52.66)} = 0.83$, $p = 0.48$ | Treatment $F_{(1, 162)} = 3.09$, $p = 0.08$ | Reward rate $F_{(1, 18)} = 1.30$, $p = 0.27$ |
| Session x Treatment $F_{(9, 162)} = 1.39$, $p = 0.20$ | Session x Reward rate $F_{(5.67, 102.0)} = 0.99$, $p = 0.44$ | Treatment x Reward rate $F_{(1, 162)} = 0.16$, $p = 0.69$ | Three-way interaction $F_{(9, 162)} = 1.04$, $p = 0.41$ |
| Session 11 Two-way mixed-effects model | Reward rate $F_{(1, 18)} = 1.14$, $p = 0.30$ | Treatment $F_{(1, 18)} = 3.34$, $p = 0.08$ | Two-way interaction $F_{(1, 18)} = 0.37$, $p = 0.55$ |
| **Panel B – Expression: Pavlovian Reward Rate task** | | | |
| Sessions 1-10 Three-way mixed-effects model | Session $F_{(2.13, 38.25)} = 19.80$, $\mathbf{p < 0.0001}$ | Treatment $F_{(1, 162)} = 0.1$, $p = 0.91$ | Reward rate $F_{(1, 18)} = 1.66$, $p = 0.21$ |
| Session x Treatment $F_{(9, 162)} = 1.19$, $p = 0.31$ | Session x Reward rate $F_{(5.42, 97.60)} = 1.95$, $p = 0.09$ | Treatment x Reward rate $F_{(1, 162)} = 0.07$, $p = 0.79$ | Three-way interaction $F_{(9, 162)} = 0.93$, $p = 0.50$ |
| Sessions 11-20 Three-way mixed-effects model | Session $F_{(3.18, 57.28)} = 1.35$, $p = 0.27$ | Treatment $F_{(1, 162)} = 15.09$, $\mathbf{p = 0.0001}$ | Reward rate $F_{(1, 18)} = 0.004$, $p = 0.95$ |
| Session x Treatment $F_{(9, 162)} = 2.42$, $\mathbf{p < 0.01}$ | Session x Reward rate $F_{(5.25, 94.41)} = 4.22$, $\mathbf{p = 0.001}$ | Treatment x Reward rate $F_{(1, 162)} = 1.63$, $p = 0.20$ | Three-way interaction $F_{(9, 162)} = 1.35$, $p = 0.22$ |
| Sessions 21-30 Three-way mixed-effects model | Session $F_{(2.76, 49.75)} = 1.14$, $p = 0.34$ | Treatment $F_{(1, 162)} = 0.79$, $p = 0.37$ | Reward rate $F_{(1, 18)} = 2.39$, $p = 0.14$ |
| Session x Treatment $F_{(9, 162)} = 1.17$, $p = 0.32$ | Session x Reward rate $F_{(4.80, 86.44)} = 0.42$, $p = 0.83$ | Treatment x Reward rate $F_{(1, 162)} = 0.01$, $p = 0.94$ | Three-way interaction $F_{(9, 162)} = 1.09$, $p = 0.38$ |
| Session 21 Two-way mixed-effects model | Reward rate $F_{(1, 18)} = 0.59$, $p = 0.45$ | Treatment $F_{(1, 18)} = 0.49$, $p = 0.49$ | Two-way interaction $F_{(1, 18)} = 0.00003$, $p = 0.99$ |

| **Table 5: Figure 5 – Post 9s US head entries – Pavlovian Reward Size task** | | | |
|---|---|---|---|
| **Panel B – Acquisition: Pavlovian Reward Size task** | | | |
| Sessions 1-10 Three-way mixed-effects model | Session $F_{(3.51, 49.15)} = 1.64$, $p = 0.19$ | Treatment $F_{(1, 126)} = 16.73$, $\mathbf{p < 0.0001}$ | Reward size $F_{(1, 14)} = 1.28$, $p = 0.28$ |
| Session x Treatment $F_{(9, 126)} = 2.11$, $\mathbf{p = 0.03}$ | Session x Reward size $F_{(3.29, 45.98)} = 2.08$, $p = 0.11$ | Treatment x Reward size $F_{(1, 126)} = 3.18$, $p = 0.08$ | Three-way interaction $F_{(9, 126)} = 2.82$, $\mathbf{p = 0.005}$ |
| Sessions 11-20 Three-way mixed-effects model | Session $F_{(1.72, 24.01)} = 1.11$, $p = 0.34$ | Treatment $F_{(1, 126)} = 3.35$, $p = 0.07$ | Reward size $F_{(1, 14)} = 3.15$, $p = 0.10$ |
| Session x Treatment $F_{(9, 126)} = 0.78$, $p = 0.64$ | Session x Reward size $F_{(4.22, 59.06)} = 1.60$, $p = 0.18$ | Treatment x Reward size $F_{(1, 126)} = 1.14$, $p = 0.29$ | Three-way interaction $F_{(9, 126)} = 0.59$, $p = 0.80$ |
| Sessions 21-30 Three-way mixed-effects model | Session $F_{(2.05, 28.75)} = 1.12$, $p = 0.34$ | Treatment $F_{(1, 126)} = 4.47$, $\mathbf{p = 0.04}$ | Reward size $F_{(1, 14)} = 9.51$, $\mathbf{p = 0.008}$ |
| Session x Treatment $F_{(9, 126)} = 1.09$, $p = 0.38$ | Session x Reward size $F_{(4.17, 58.44)} = 1.79$, $p = 0.14$ | Treatment x Reward size $F_{(1, 126)} = 2.17$, $p = 0.14$ | Three-way interaction $F_{(9, 126)} = 1.08$, $p = 0.38$ |
| Session 11 Two-way mixed-effects model | Reward size $F_{(1, 14)} = 2.49$, $p = 0.14$ | Treatment $F_{(1, 14)} = 5.48$, $\mathbf{p = 0.03}$ | Two-way interaction $F_{(1, 14)} = 2.29$, $p = 0.15$ |
| **Panel C – Expression: Pavlovian Reward Size task** | | | |
| Sessions 1-10 Three-way mixed-effects model | Session $F_{(3.68, 51.54)} = 0.43$, $p = 0.77$ | Treatment $F_{(1, 126)} = 1.55$, $p = 0.22$ | Reward size $F_{(1, 14)} = 6.21$, $\mathbf{p = 0.03}$ |
| Session x Treatment $F_{(9, 126)} = 1.70$, $p = 0.09$ | Session x Reward size $F_{(3.49, 48.80)} = 2.98$, $\mathbf{p = 0.03}$ | Treatment x Reward size $F_{(1, 126)} = 0.01$, $p = 0.92$ | Three-way interaction $F_{(9, 126)} = 1.95$, $p = 0.05$ |
| Sessions 11-20 Three-way mixed-effects model | Session $F_{(2.00, 28.06)} = 0.52$, $p = 0.60$ | Treatment $F_{(1, 126)} = 5.23$, $\mathbf{p = 0.02}$ | Reward size $F_{(1, 14)} = 2.30$, $p = 0.15$ |
| Session x Treatment $F_{(9, 126)} = 1.29$, $p = 0.25$ | Session x Reward size $F_{(4.77, 66.71)} = 0.56$, $p = 0.73$ | Treatment x Reward size $F_{(1, 126)} = 1.62$, $p = 0.21$ | Three-way interaction $F_{(9, 126)} = 0.84$, $p = 0.58$ |
| Sessions 21-30 Three-way mixed-effects model | Session $F_{(1.89, 26.47)} = 1.23$, $p = 0.31$ | Treatment $F_{(1, 126)} = 4.61$, $\mathbf{p = 0.03}$ | Reward size $F_{(1, 14)} = 7.34$, $\mathbf{p = 0.02}$ |
| Session x Treatment $F_{(9, 126)} = 1.66$, $p = 0.11$ | Session x Reward size $F_{(4.05, 56.72)} = 0.55$, $p = 0.70$ | Treatment x Reward size $F_{(1, 126)} = 3.35$, $p = 0.07$ | Three-way interaction $F_{(9, 126)} = 1.36$, $p = 0.22$ |
| Session 21 Two-way mixed-effects model | Reward size $F_{(1, 14)} = 4.11$, $p = 0.06$ | Treatment $F_{(1, 14)} = 1.82$, $p = 0.20$ | Two-way interaction $F_{(1, 14)} = 3.56$, $p = 0.08$ |

| Table 6: Figure 6 – Post 9s US head entries – Pavlovian Reward Rate task | | | |
|---|---|---|---|
| **Panel B – Acquisition: Pavlovian Reward Rate task** | | | |
| Sessions 1-10<br>Three-way mixed-effects model | Session<br>$F_{(4.23, 76.09)} = 6.57$, $\boldsymbol{p = 0.0001}$ | Treatment<br>$F_{(1, 162)} = 4.29$, $\boldsymbol{p = 0.04}$ | Reward rate<br>$F_{(1, 18)} = 0.05$, $p = 0.83$ |
| Session x Treatment<br>$F_{(9, 162)} = 1.09$, $p = 0.37$ | Session x Reward rate<br>$F_{(4.40, 79.21)} = 0.28$, $p = 0.91$ | Treatment x Reward rate<br>$F_{(1, 162)} = 0.22$, $p = 0.64$ | Three-way interaction<br>$F_{(9, 162)} = 0.74$, $p = 0.67$ |
| Sessions 11-20<br>Three-way mixed-effects model | Session<br>$F_{(5.09, 91.55)} = 1.45$, $p = 0.21$ | Treatment<br>$F_{(1, 162)} = 2.28$, $p = 0.13$ | Reward rate<br>$F_{(1, 18)} = 0.09$, $p = 0.77$ |
| Session x Treatment<br>$F_{(9, 162)} = 1.12$, $p = 0.35$ | Session x Reward rate<br>$F_{(4.80, 86.41)} = 0.86$, $p = 0.51$ | Treatment x Reward rate<br>$F_{(1, 162)} = 0.08$, $p = 0.77$ | Three-way interaction<br>$F_{(9, 162)} = 1.28$, $p = 0.25$ |
| Sessions 21-30<br>Three-way mixed-effects model | Session<br>$F_{(4.96, 89.20)} = 0.86$, $p = 0.51$ | Treatment<br>$F_{(1, 162)} = 2.82$, $p = 0.10$ | Reward rate<br>$F_{(1, 18)} = 0.68$, $p = 0.42$ |
| Session x Treatment<br>$F_{(9, 162)} = 1.07$, $p = 0.39$ | Session x Reward rate<br>$F_{(4.28, 76.98)} = 1.87$, $p = 0.12$ | Treatment x Reward rate<br>$F_{(1, 162)} = 0.03$, $p = 0.86$ | Three-way interaction<br>$F_{(9, 162)} = 1.58$, $p = 0.13$ |
| Session 11<br>Two-way mixed-effects model | Reward rate<br>$F_{(1, 18)} = 0.42$, $p = 0.52$ | Treatment<br>$F_{(1, 18)} = 1.89$, $p = 0.19$ | Two-way interaction<br>$F_{(1, 18)} = 1.23$, $p = 0.28$ |
| **Panel C – Expression: Pavlovian Reward Rate task** | | | |
| Sessions 1-10<br>Three-way mixed-effects model | Session<br>$F_{(4.65, 83.67)} = 8.32$, $\boldsymbol{p < 0.0001}$ | Treatment<br>$F_{(1, 162)} = 0.01$, $p = 0.92$ | Reward rate<br>$F_{(1, 18)} = 0.25$, $p = 0.63$ |
| Session x Treatment<br>$F_{(9, 162)} = 0.87$, $p = 0.56$ | Session x Reward rate<br>$F_{(4.90, 88.16)} = 0.15$, $p = 0.70$ | Treatment x Reward rate<br>$F_{(1, 162)} = 1.26$, $p = 0.26$ | Three-way interaction<br>$F_{(9, 162)} = 1.85$, $p = 0.06$ |
| Sessions 11-20<br>Three-way mixed-effects model | Session<br>$F_{(3.05, 54.84)} = 1.39$, $p = 0.26$ | Treatment<br>$F_{(1, 162)} = 0.64$, $p = 0.42$ | Reward rate<br>$F_{(1, 18)} = 1.04$, $p = 0.32$ |
| Session x Treatment<br>$F_{(9, 162)} = 1.88$, $p = 0.06$ | Session x Reward rate<br>$F_{(3.79, 68.15)} = 1.57$, $p = 0.19$ | Treatment x Reward rate<br>$F_{(1, 162)} = 3.22$, $p = 0.07$ | Three-way interaction<br>$F_{(9, 162)} = 0.79$, $p = 0.62$ |
| Sessions 21-30<br>Three-way mixed-effects model | Session<br>$F_{(5.10, 91.81)} = 0.78$, $p = 0.57$ | Treatment<br>$F_{(1, 162)} = 0.38$, $p = 0.54$ | Reward rate<br>$F_{(1, 18)} = 0.97$, $p = 0.34$ |
| Session x Treatment<br>$F_{(9, 162)} = 0.35$, $p = 0.96$ | Session x Reward rate<br>$F_{(4.49, 80.75)} = 1.10$, $p = 0.37$ | Treatment x Reward rate<br>$F_{(1, 162)} = 3.06$, $p = 0.06$ | Three-way interaction<br>$F_{(9, 162)} = 0.69$, $p = 0.72$ |
| Session 21<br>Two-way mixed-effects model | Reward rate<br>$F_{(1, 18)} = 0.0002$, $p = 0.99$ | Treatment<br>$F_{(1, 18)} = 1.57$, $p = 0.23$ | Two-way interaction<br>$F_{(1, 18)} = 0.64$, $p = 0.43$ |

| Table 7: Fig 7 – Total head entries | | | |
|---|---|---|---|
| **Panel A – Acquisition: Pavlovian Reward Size task** | | | |
| Sessions 1-10<br>Two-way mixed-effects model | Session<br>$F_{(9, 126)} = 2.24$, $\boldsymbol{p = 0.02}$ | Treatment<br>$F_{(1, 14)} = 16.24$, $\boldsymbol{p = 0.001}$ | Session x Treatment<br>$F_{(9, 126)} = 1.58$, $p = 0.13$ |
| Sessions 11-20<br>Two-way mixed-effects model | Session<br>$F_{(9, 126)} = 1.43$, $p = 0.18$ | Treatment<br>$F_{(1, 14)} = 2.54$, $p = 0.13$ | Session x Treatment<br>$F_{(9, 126)} = 1.50$, $p = 0.15$ |
| Sessions 21-30<br>Two-way mixed-effects model | Session<br>$F_{(9, 126)} = 1.97$, $\boldsymbol{p = 0.05}$ | Treatment<br>$F_{(1, 14)} = 0.98$, $p = 0.34$ | Session x Treatment<br>$F_{(9, 126)} = 5.51$, $\boldsymbol{p < 0.0001}$ |
| **Panel A – Expression: Pavlovian Reward Size task** | | | |
| Sessions 1-10<br>Two-way mixed-effects model | Session<br>$F_{(9, 126)} = 3.33$, $\boldsymbol{p = 0.001}$ | Treatment<br>$F_{(1, 14)} = 0.0004$, $p = 0.98$ | Session x Treatment<br>$F_{(9, 126)} = 0.95$, $p = 0.48$ |
| Sessions 11-20<br>Two-way mixed-effects model | Session<br>$F_{(9, 126)} = 1.10$, $p = 0.36$ | Treatment<br>$F_{(1, 14)} = 27.09$, $\boldsymbol{p = 0.001}$ | Session x Treatment<br>$F_{(9, 126)} = 2.08$, $\boldsymbol{p = 0.04}$ |
| Sessions 21-30<br>Two-way mixed-effects model | Session<br>$F_{(9, 126)} = 0.78$, $p = 0.63$ | Treatment<br>$F_{(1, 14)} = 6.89$, $\boldsymbol{p = 0.02}$ | Session x Treatment<br>$F_{(9, 126)} = 1.75$, $p = 0.09$ |
| **Panel B – Acquisition: Pavlovian Reward Rate task** | | | |
| Sessions 1-10<br>Two-way mixed-effects model | Session<br>$F_{(9, 162)} = 0.76$, $p = 0.65$ | Treatment<br>$F_{(1, 18)} = 9.47$, $\boldsymbol{p = 0.007}$ | Session x Treatment<br>$F_{(9, 162)} = 3.98$, $\boldsymbol{p = 0.0001}$ |
| Sessions 11-20<br>Two-way mixed-effects model | Session<br>$F_{(9, 162)} = 0.91$, $p = 0.52$ | Treatment<br>$F_{(1, 18)} = 3.02$, $p = 0.10$ | Session x Treatment<br>$F_{(9, 162)} = 1.04$, $p = 0.41$ |
| Sessions 21-30<br>Two-way mixed-effects model | Session<br>$F_{(9, 162)} = 2.90$, $\boldsymbol{p = 0.003}$ | Treatment<br>$F_{(1, 18)} = 1.76$, $p = 0.20$ | Session x Treatment<br>$F_{(9, 162)} = 2.25$, $\boldsymbol{p = 0.02}$ |

| **Panel B – Expression: Pavlovian Reward Rate task** | | | |
|---|---|---|---|
| Sessions 1-10<br>Two-way mixed-effects model | Session<br>$F_{(9, 162)} = 1.27$, $p = 0.26$ | Treatment<br>$F_{(1, 18)} = 1.58$, $p = 0.23$ | Session x Treatment<br>$F_{(9, 162)} = 0.25$, $p = 0.25$ |
| Sessions 11-20<br>Two-way mixed-effects model | Session<br>$F_{(9, 162)} = 1.59$, $p = 0.12$ | Treatment<br>$F_{(1, 18)} = 11.85$, $\boldsymbol{p} \mathbf{= 0.003}$ | Session x Treatment<br>$F_{(9, 162)} = 1.27$, $p = 0.26$ |
| Sessions 21-30<br>Two-way mixed-effects model | Session<br>$F_{(9, 162)} = 2.21$, $\boldsymbol{p} \mathbf{= 0.02}$ | Treatment<br>$F_{(1, 18)} = 0.03$, $p = 0.86$ | Session x Treatment<br>$F_{(9, 162)} = 1.66$, $p = 0.10$ |
